# Minute amounts of helicase-deficient truncated RECQL4 are sufficient for DNA replication

**DOI:** 10.1101/2025.07.21.666025

**Authors:** Paula Armina V. Buco, Wilson Tandazo-Castillo, Alistair M. Chalk, Courtney Pilcher, Jessica K. Holien, Jörg Heierhorst, Tiong Y. Tan, Amnon Koren, Monique F. Smeets, Carl R. Walkley

## Abstract

RECQL4 is a member of the RecQ family of helicases, playing essential roles in DNA replication and maintaining genome integrity. Mutations in RECQL4 are linked to severe human diseases, including Rothmund-Thomson Syndrome, RAPIDALINO Syndrome, and Baller-Gerold Syndrome. However, we still do not fully understand its functions and genetic interactions. The role of the ATP-dependent helicase activity in RECQL4 remains controversial. To understand RECQL4’s functions further, we conducted a genome-wide forward genetic screen using murine models that closely mimic the RECQL4 mutations found in patients with Rothmund-Thomson syndrome. Our goal was to identify loss-of-function alleles that could rescue the proliferation and viability defects associated with RECQL4 mutation. From our screening we identified the loss of KLHDC3, a substrate-binding subunit of the Cullin-RING ligase (CRL) E3, as the most significant rescue allele. KLHDC3 facilitates the ubiquitin-mediated destruction of proteins with specific C-terminal degron motifs. Its loss normalized cell proliferation and DNA replication rates in cells with mutated RECQL4. Further analysis revealed that the loss of KLHDC3 led to the stabilization of minute levels of a truncated RECQL4 protein. This RECQL4 fragment contained a neo-degron sequence specific for KLHDC3, formed after Cre-mediated recombination of the *Recql4^fl^* allele. Although this rescue mechanism does not apply to human RECQL4 mutations, it shows that very low chromatin-bound levels of a truncated RECQL4 protein—comprising only the N-terminal 480 amino acids, including its Sld2-like domain but lacking the ATP-dependent helicase domain and the entire C-terminal portion—are sufficient to support DNA replication in mammalian cells. These results demonstrate that the ATPase activity and helicase domain of RECQL4 are not essential for DNA replication in mammals. Furthermore, our findings suggest that there are unlikely to be monogenic loss-of-function alleles that can rescue RECQL4 mutations. This demonstrates that RECQL4 is an essential and non-redundant regulator of DNA replication and cell viability and that this activity does not require the ATP dependent helicase activity.

## Introduction

The orderly and accurate duplication of DNA during each cell division cycle is essential for development and homeostasis of all organisms. Despite the fundamental importance of DNA replication, there are significant knowledge gaps about the specific roles of numerous replication proteins. One such protein is RecQ like helicase 4 (RECQL4). RECQL4 is essential for DNA replication across multicellular organisms from Drosophila to mammals (Chu & Hickson, 2009).

Although RECQL4 was long considered the mammalian homologue of the essential yeast DNA replication factor Sld2, it is now clear that a structurally unrelated protein DONSON has evolved to carry out the Sld2-like function during mammalian DNA replication (Cvetkovic *et al*, 2023; Evrin *et al*, 2023; Hashimoto *et al*, 2023; Kingsley *et al*, 2023; Lim *et al*, 2023). As a result, the role and function of RECQL4 in normal DNA replication remains a major knowledge gap in our understanding of this fundamental cellular process. This knowledge will be directly relevant to understanding human disease, as mutations in RECQL4, like those of the other mammalian RecQ helicases BLM, WRN and RECQL5, are associated with human disease (Chu & Hickson, 2009; Hickson, 2003; Lu & Davis, 2021).

RECQL4 is a member of the RecQ family of helicases with genome integrity functions, similar to the enzymes mutated in Bloom and Werner syndromes (Hickson, 2003; Kitao *et al*, 1999a; Kitao *et al*, 1999b; Wang *et al*, 2003; Wang *et al*, 2001; Wang *et al*, 2002). Bi-allelic compound heterozygous mutations in *RECQL4* have been reported in three rare autosomal recessive human genetic syndromes: RAPADILINO Syndrome (OMIM #266280), Baller-Gerold Syndrome (BGS, OMIM #218600) and Rothmund-Thomson Syndrome (RTS, OMIM #268400). A significant body of work has reported that RECQL4 is required for DNA replication (Sangrithi *et al*, 2005) and is involved in various DNA damage repair pathways (Fielden *et al*, 2025; Jin *et al*, 2008; Lu *et al*, 2017; Petkovic *et al*, 2005; Singh *et al*, 2010; Thakur *et al*, 2025). Mouse models that lack RECQL4 protein expression led to early-to-mid-gestational lethality (Ichikawa *et al*, 2002; Lu *et al*, 2015; Smeets *et al*, 2014). In contrast, mice with a targeted biochemical mutation that specifically abolished ATP dependent helicase activity (murine p.K525A, homologous to human p.K508A) were born at Mendelian frequency, fertile, not spontaneously cancer prone and had a normal lifespan under standard housing conditions (Castillo-Tandazo *et al*, 2019). This result demonstrated that the ATP-dependent helicase activity of RECQL4 was not essential *in vivo* for DNA replication or murine homeostasis, consistent with analysis in human cell line models (Padayachy *et al*, 2024).

In contrast to these findings that RECQL4’s role in DNA replication is independent of its ATPase activity, a recent single molecule analysis in reconstituted Xenopus egg extracts proposed that the helicase activity of RECQL4 was required to evict DONSON from the Cdc45/Mcm2-7/GINS (CMG) complex to allow DNA replication (Terui *et al*, 2024). The authors of this recent study noted that whilst they considered that they had successfully depleted endogenous xRECQL4 from the extracts, the extract still retained some origin firing activity potentially indicative of trace amounts of RECQL4 remaining (Terui *et al*., 2024). At present it is not clear how these seemingly contradictory findings from mouse models and human cell lines can be reconciled with the recent single molecule analysis in Xenopus.

Herein we have used forward genetics in murine cells to identify suppressor mutations that rescued DNA replication in the presence of RECQL4 mutation. This approach led to the identification of KLHDC3, the substrate-binding subunit of the Cullin-RING ligase (CRL) E3 that facilitates ubiquitin-mediated destruction of proteins with specific C-terminal degron motifs (Pilcher *et al*, 2025; Scott *et al*, 2024a). Loss of KLHDC3 normalised cell proliferation and DNA replication rates in *Recql4* mutated cells. This occurred by KLHDC3 loss leading to stabilisation of trace levels of a truncated RECQL4 protein containing a neo-degron specific for KLHDC3, formed after Cre-mediated recombination of the *Recql4^fl^*allele. Whilst this rescue mechanism is restricted to this specific model and not generally applicable to RECQL4 mutation, it demonstrates that very low levels of truncated RECQL4 - containing only the N-terminal 480 amino acids in-frame including its Sld2-like domain but lacking the ATP-dependent helicase domain and entire C-terminal portion of the protein – was capable of supporting DNA replication *in vivo* in mammalian cells. These results provide an orthogonal system to our previous work with a knock-in allele of a ATP binding mutant (Castillo-Tandazo *et al*., 2019) demonstrating that the ATPase activity and helicase domain of RECQL4 are not essential for mammalian DNA replication.

## Results

### Genome-wide loss of function screen to identify suppressors of RECQL4 mutation phenotypes

Our previous work using an allelic series of *Recql4* mutant mouse models had identified that *in vivo* loss-of-function RECQL4 mutations resulted in a fully penetrant bone marrow (BM) failure phenotype (Castillo-Tandazo *et al*., 2021; Castillo-Tandazo *et al*., 2019; Smeets *et al*., 2014). This did not occur in mice engineered to have an ATPase dead RECQL4 either as a germ-line mutation (*Recql4^K525A/K525A^*) or upon acute expression (*R26*-CreER^T2^ *Recql4^1/K525A^*) (Castillo-Tandazo *et al*., 2019), indicating that the BM failure was not related to RECQL4’s ATP dependent helicase function. In contrast, mouse models acutely expressing a truncated but mislocalised (p.R347X) or truncated and unstable (p.G522EfsX6) RECQL4 protein developed BM failure (Castillo-Tandazo *et al*., 2021; Castillo-Tandazo *et al*., 2019). The p.R347X lacks the Zn knuckle domain and most of the region required for ssDNA and fork DNA binding and has altered localisation (Castillo-Tandazo *et al*., 2021; Castillo-Tandazo *et al*., 2019). The p.G522Efs product is unstable and we have not been able to detect any protein product either of native protein or when GFP-fused and expressed as a cDNA (Castillo-Tandazo *et al*., 2021; Castillo-Tandazo *et al*., 2019).

To establish a tractable model suitable for studying this phenomenon we generated immortalised myeloid cell lines from the BM of the different *Recql4* alleles crossed to the *Rosa26-* CreER^T2^ allele (Castillo-Tandazo *et al*., 2019; Heraud-Farlow *et al*., 2024; Wang *et al*., 2006; Xu *et al*., 2022). In these cells, the floxed *Recql4^fl^* allele is deleted upon tamoxifen-induced activation of the CreER^T2^ recombinase. We observed that a homozygous null allele (*Recql4^fl/fl^*) of RECQL4 resulted in rapid proliferation arrest and cell death upon Cre activation compared to *Recql4^fl/+^* heterozygous cells (that retained one WT allele) used as a control. However, we also observed that the floxed allele deletion in the *Recql4^R347X/Δ^* retained a certain level of proliferation and viability (Castillo-Tandazo *et al*., 2021; Ng *et al*., 2015). We therefore chose to perform a screen using the p.R347X allele.

To identify genes whose loss restored proliferation and viability to *Recql4* mutant cell lines, we engineered *R26*CreER^T2^ *Recql4^fl/+^* and *R26*CreER^T2^ *Recql4^fl/R347X^* cell lines to constitutively express Cas9 for a loss-of-function genome-wide screen using a murine Brie sgRNA library (Figure 1A) (Doench *et al*., 2016; Heraud-Farlow *et al*., 2024; Xu *et al*., 2022). After selection of Brie library infected cells, tamoxifen was added to delete the floxed *Recql4* allele (day 0). The cells were then cultured with tamoxifen, counted and their DNA pellets collected at day 0, day 4 and day 9 and the sgRNA copy number was determined by sequencing (Figure 1A-1B). The sgRNA copy number was compared between Days 4 and 9 to Day 0, respectively (Figure S1A-S1F). Surprisingly, we identified only a single candidate that was significantly enriched in the *Recql4^1/R347X^* cells compared to the *Recql4^+/Δ^* control cells, sgRNA against Kelch domain containing 3 (*Klhdc3*) (Figure 1B; Dataset S1).

**Figure 1.**
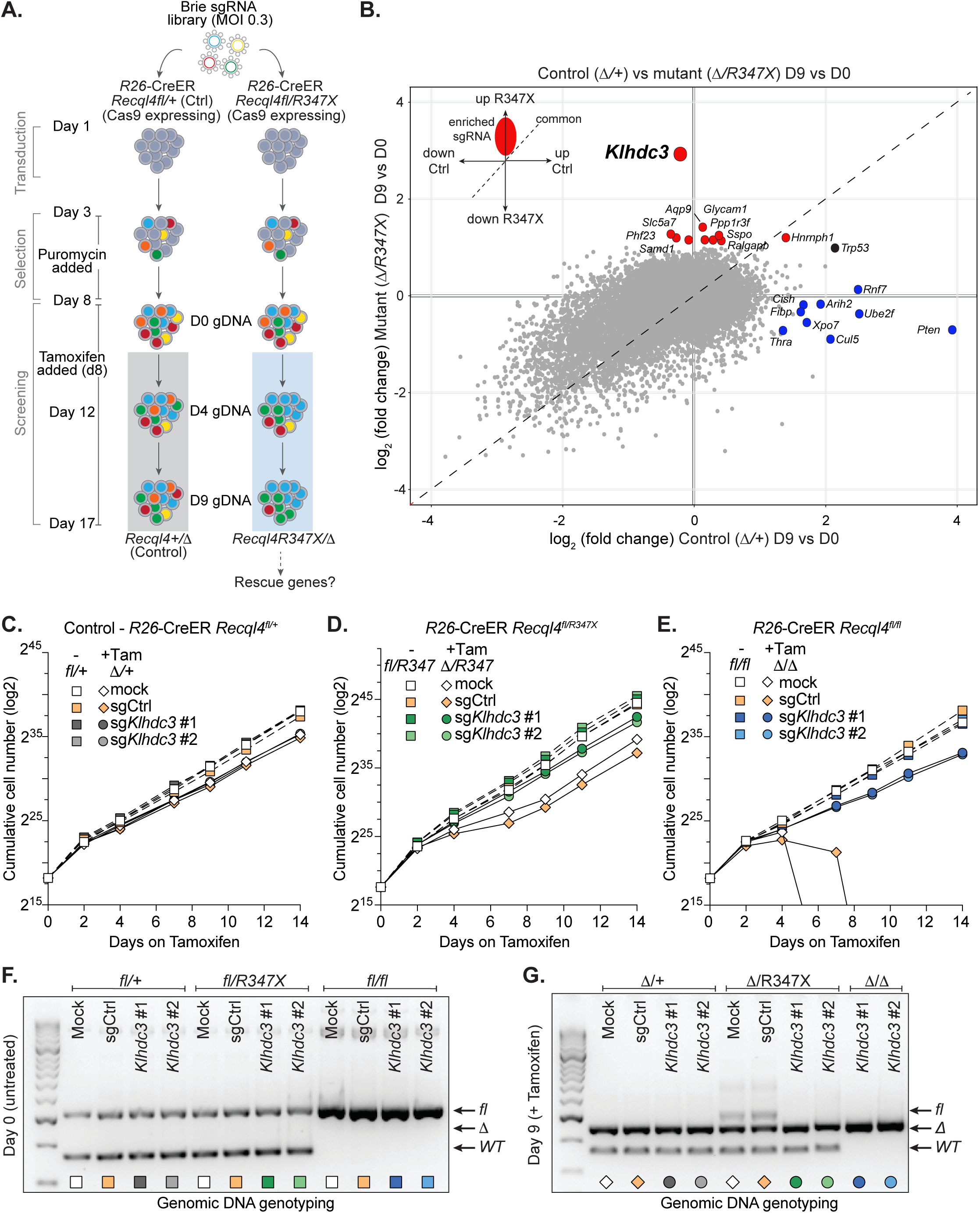
Loss of *Klhdc3* rescues *Recql4* loss-of-function mutant myeloid cells. A. Schematic outline of genetic screen designed to identify loss-of-function rescue alleles of proliferation defects induced by RECQL4 p.R347X mutation. B. Summary of MaGeCK maximum likelihood estimation (mle) analysis of screen showing enrichment of sgRNAs targeting the indicated genes in the Δ*/R347X* cells compared with those in the Δ*/+* cells at day 9/day 0. Red dots, high-confidence indicate statistically significant enrichment in the Δ*/R347X* cells; blue indicated enriched sgRNA in the Δ*/+* cells. C. Effect of loss of *Klhdc3* on control cells (*R26*-CreER *Recql4^fl/+^*; become Δ*/+* cell following tamoxifen treatment applied at Day 0) proliferation. D. Effect of loss of *Klhdc3* on RECQL4 p.R347X only expressing cells (*R26*-CreER *Recql4^fl/R347X^*; become Δ*/R347X* cells following tamoxifen treatment applied at Day 0) proliferation. E. Effect of loss of *Klhdc3* on Recql4 deficient cells (*R26*-CreER *Recql4^fl/fl^*; become Δ*/*Δ cells following tamoxifen treatment applied at Day 0) proliferation. F. Genotyping PCR using genomic DNA of the *Recql4* locus at Day 0; each allele as indicated. G. Genotyping PCR using genomic DNA of the *Recql4* locus at Day 9 of tamoxifen treatment; each allele as indicated.

KLHDC3 is a substrate receptor of the Cullin2-RING ligase (CRL2) that recognises and facilitates ubiquitin-mediated destruction of proteins with specific C-terminal motifs via the recently discovered C-end degron (DesCEND) pathway (Koren *et al*, 2018; Lin *et al*., 2018; Scott *et al*., 2024a; Scott *et al*, 2024b; Timms *et al*, 2023; Zhang *et al*, 2023). Assessment of the screen data revealed no enrichment of loss-of-function alleles of any other KLHDC family member nor other proteins known to interact with KLHDC3 (such as the other components of the CRL E3 complex), indicating that the mechanism of rescue was specific to loss of KLHDC3.

Loss of KLHDC3 did not have an appreciable effect on the control cells (*Recql4* heterozygous; fl/+) and the sg*Klhdc3* guides were not enriched in the control genotype (Figure 1B). The loss of KLHDC3 was first validated in the same two Cas9 cell lines (Fig S2D-F) and then in three additional (non-Cas9 expressing) cell lines including a *Recql4^fl/fl^*cell line (Fig 1C-G; S2A-C). We validated the result using 4 different sgRNA (the top 2 ranked sgRNA from the Brie library and 2 additional sgRNA not present in the library) targeting different regions of *Klhdc3* in *R26*-CreER^T2^ *Recql4^fl/R347X^* cell lines (Figure 1C-D; Figure S2A-2B, S2D). To exclude survival and outgrowth of *R26*CreER^T2^ *Recql4^fl/R347X^*cells that had not fully recombined the *Recql4^fl^* allele, we completed genotyping of the deletion efficiency in sgControl (sgCtrl) and sg*Klhdc3* cells. Strikingly, in sg*Klhdc3* cells there was complete and stable recombination of the *Recql4^fl^* allele with complete loss of the full-length WT protein and stable expression of the p.R347X at ∼28% of WT levels (Figure 1F-1G; Figure S2E-S2F). Sequencing of the DNA confirmed this and that the *Klhdc3* mutation was homozygous and predicted to be deleterious (across all sgRNA used for *Klhdc3*) (Figure S3A-S3C). The *Recql4^1/R347X^* sg*Klhdc3* cells (referred to hereon as *Recql4^1/R347X^ Klhdc3^1/1^* cells) had a proliferation rate comparable to tamoxifen treated control cells irrespective of the *Recql4* allele status.

These results prompted us to test if cells predicted to be null for RECQL4 (*R26*-CreER^T2^ *Recql4^fl/fl^*) would also be rescued by loss of KLHDC3. To our surprise, given the profound and rapid proliferation failure and cell death engendered by tamoxifen treatment of the *R26*-CreER^T2^ *Recql4^fl/fl^* cells (Figure 1E), the removal of KLHDC3 prior to tamoxifen treatment enabled the *Recql4^1/1^ Klhdc3^1/1^* cells to survive and proliferate at the same rate as control cells (Figure 1E-1G; Figure S2C).

Therefore, this forward genetic screen identified that the loss of KLHDC3 specifically and uniquely allowed for sustained proliferation and viability of RECQL4 deficient or mutant expressing myeloid cells.

### Enhanced DNA replication licensing in the *Recql4^1/1^ Klhdc3^1/1^* cells

We next sought to understand the characteristics of the rescued *Recql4^1/1^ Klhdc3^1/1^* cells. We isolated and validated clones from two independently derived cell lines (made from different donor mouse bone marrows). These were confirmed as *Klhdc3* homozygous mutant and had undergone complete genomic recombination of the floxed *Recql4* allele with no RECQL4 protein detectable in whole cell western blot (Figure 1G; Figure S2E, S2G). Analysis of proliferation rates showed that the *Recql4^1/1^ Klhdc3^1/1^* cells had a comparable proliferation rate to control cells, including *Klhdc3^1/1^* single mutant cells (Figure 1C; Figure S2A). We then assessed DNA replication rates using pulse labelling of asynchronous cultures of myeloid cells with 5-ethynyl-2′-deoxyuridine (EdU) incorporation (Figure 2A). The EdU pulse labelling demonstrated that sg*Klhdc3* cells were indistinguishable from sgCtrl myeloid cells, indicating that loss of KLHDC3 alone did not change DNA replication and cell cycle phase entry rates perceptibly (Figure 2A). Interestingly, whilst the overall cell cycle phase distribution of the *Recql4^1/1^* sg*Klhdc3* was not significantly different from either sgCtrl or sg*Klhdc3*, there was a reproducible reduction in the overall intensity of the EdU signal in S phase of the cell cycle (Figure 2A). This suggested a lower overall rate of EdU incorporation in the *Recql4^1/1^ Klhdc3^1/1^* cells, with a more prominent reduction during the second half of S phase. A similar profile was also obtained upon deletion of *Recql4* in *Klhdc3* WT cells, although these cells fail to proliferate and die within 5-14 days (Figure S4A-S4B). To further confirm this change was due to the loss of RECQL4 and not an effect unique to the combination of *Recql4^1/1^ Klhdc3^1/1^* loss, we re-expressed mCherry-mRECQL4 in the rescued cells and repeated the analysis (Figure 2B). Re-expression of RECQL4 restored the EdU profiles and intensity/level of EdU signal in S phase to that of control cells. This demonstrates that the reduced EdU incorporation during S phase, despite a normal overall cell cycle phase distribution and entry time, was a specific result of RECQL4 loss (Figure 2B). To gain more insight, we coupled EdU and Ki67 staining to obtain a more fine-grained analysis of cell cycle state and DNA replication rates (Figure 2C). When compared to control cell lines, the *Recql4^1/1^ Klhdc3^1/1^* cells had evidence of continuing DNA synthesis (EdU incorporation) during mitosis (Figure 2C). Taken together, these results indicate that the *Recql4^1/1^ Klhdc3^1/1^* cells have a level of replication stress, albeit seemingly tolerated, with incomplete DNA replication during S phase, but then utilise the Mitotic DNA Synthesis pathway (MiDAS) to complete replication and sustain largely normal proliferation and viability (Bhowmick *et al*, 2023; Bhowmick *et al*, 2016; Minocherhomji *et al*, 2015).

**Figure 2.**
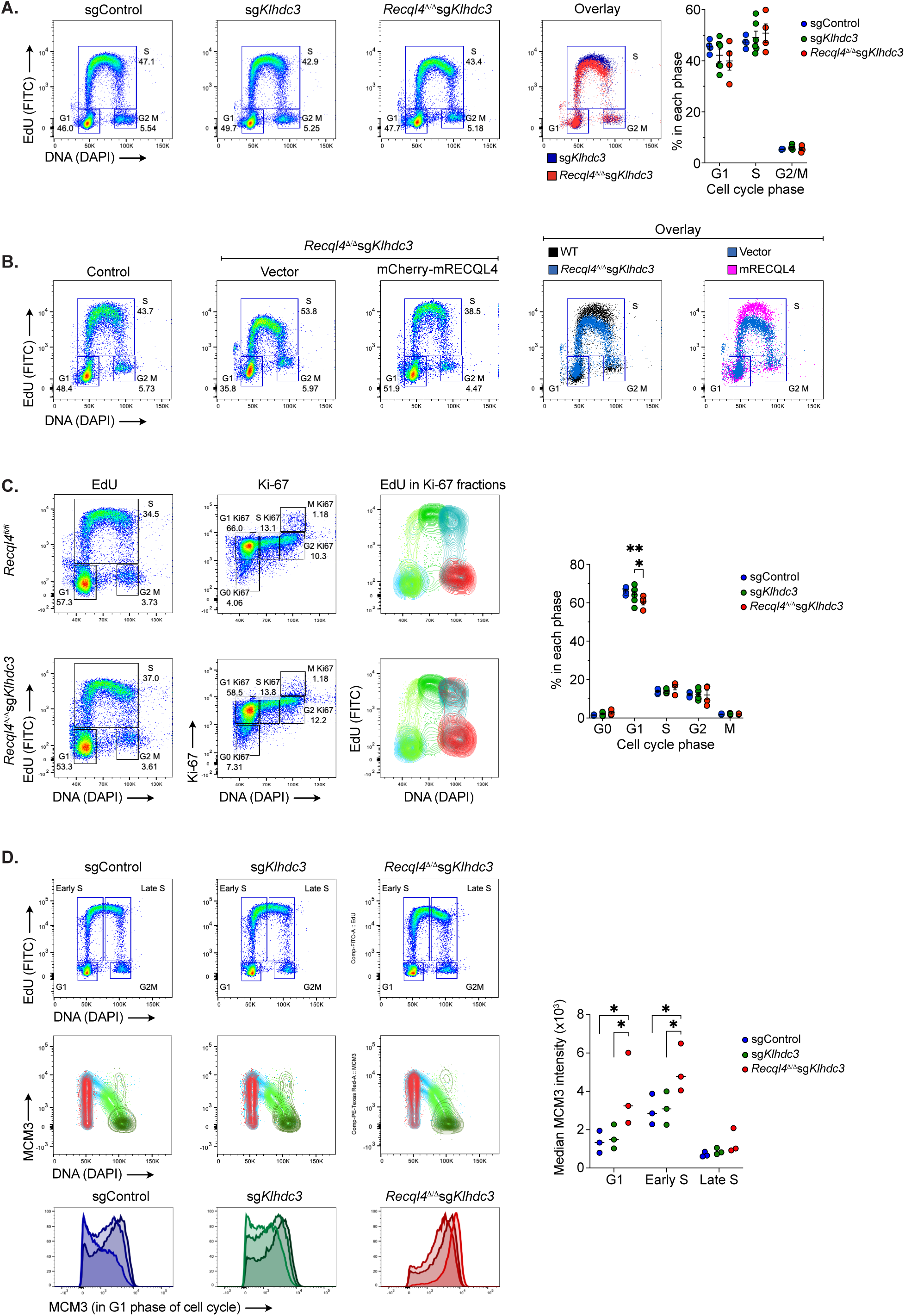
Enhanced DNA replication licensing in the *Recql4^1/1^ Klhdc3^1/1^* cells. A. Representative flow cytometry plots assessing DNA replication rates following pulse labelling of asynchronous cultures of myeloid cells with EdU incorporation and DAPI (DNA stain). Genotypes assessed as sgControl (WT); sg*Klhdc3* and *Recql4^1/1^* sg*Klhdc3*. *Recql4^1/1^* sg*Klhdc3* reproducibly had a lower maximum intensity of EdU staining. Quantitation of the cell cycle distribution from independent replicates shown. B. Re-expression of mCherry-RECQL4 protein in the *Recql4^1/1^* sg*Klhdc3* cells. Control is sgControl (WT); vector is *Recql4^1/1^* sg*Klhdc3* reconstituted with empty vector and *Recql4^1/1^* sg*Klhdc3* mCherry-RECQL4 is cells reconstituted with full length mouse RECQL4 protein. Overlays of the Edu/DAPI plots are provided. C. Analysis of cell cycle distribution using combined EdU, DAPI and Ki-67 staining. Representative flow cytometry plots for control (*Recql4^fl/fl^*; non-tamoxifen treated cells retaining *Recql4*) and *Recql4^1/1^* sg*Klhdc3* cells. First panel shows EdU vs DAPI, second panel shows Ki-67 vs DAPI and third panel shows the distribution of EdU within the fractions identified by Ki-67 staining. The *Recql4^1/1^* sg*Klhdc3* cells show a persistence of DNA replication in the G2 and M phase of the cell cycle. Quantitation of the cell cycle phases provided. D. Analysis of chromatin bound MCM3 in sgControl (WT); sg*Klhdc3* and *Recql4^1/1^* sg*Klhdc3* cells. Populations were fractionated based on cell cycle distribution based on Edu/DAPI into G1, Early S, Late S and G2/M phases as indicated (top row). The levels of chromatin loaded MCM3 were then assessed in each phase (middle row) and then specifically in the G1 phase (bottom row). The median MCM3 loading in G1 (based on fluorescence intensity) of three cell lines each genotype is shown as histogram overlays. Data expressed as mean ± sem from independent experiments; n≥3 except panel B where representative FACS plots are shown; \**P <* 0.05, \*\**P <* 0.01 as indicated with statistical comparisons from ANOVA with multiple-comparisons correction calculated using Prism software.

Based on these results we used a FACS-based method to measure the amount of the replication origin licensing factor MCM3 bound to DNA during the different cell cycle phases (Figure 2D) (Clijsters *et al*, 2019; Matson & Cook, 2020; Matson *et al*., 2017). From this analysis we found that loss of KLHDC3 alone did not change the level of MCM3 loading compared to WT cells (Figure 2D). In contrast, the *Recql4^1/1^ Klhdc3^1/1^* cells showed increased MCM3 loading intensity during G1 and early S phase compared to control cells (Figure 2D). In conclusion, we have identified that loss of KLHDC3 specifically and uniquely rescued the cellular effects of the loss or mutation of the core essential protein RECQL4. This was associated with a mild DNA replication stress and enhanced replication licensing, allowing cells to maintain an overall proliferation and viability rate comparable to RECQL4 WT cells.

### Loss of *Klhdc3 in vivo* protects against bone marrow failure induced by loss of *Recql4*

Next, we sought to determine if the rescue observed in the immortalised myeloid cells extended to *in vivo* phenotypes. To test this, we generated two *Klhdc3* mouse models: a *Klhdc3^fl/fl^* allele (Figure 3A) and a germ-line deficient *Klhdc3^+/-^* (phenotype described in detail elsewhere). These were generated on a C57BL6/J background using endonuclease mediated recombination. Both models result in the deletion of exon 4 and 5, with the germ-line deficient (*Klhdc3^+/-^*) allele arising as a result of recombination and deletion of the region encompassing the loxP flanked region during targeting. We tested *in vivo* rescue in two contexts.

**Figure 3.**
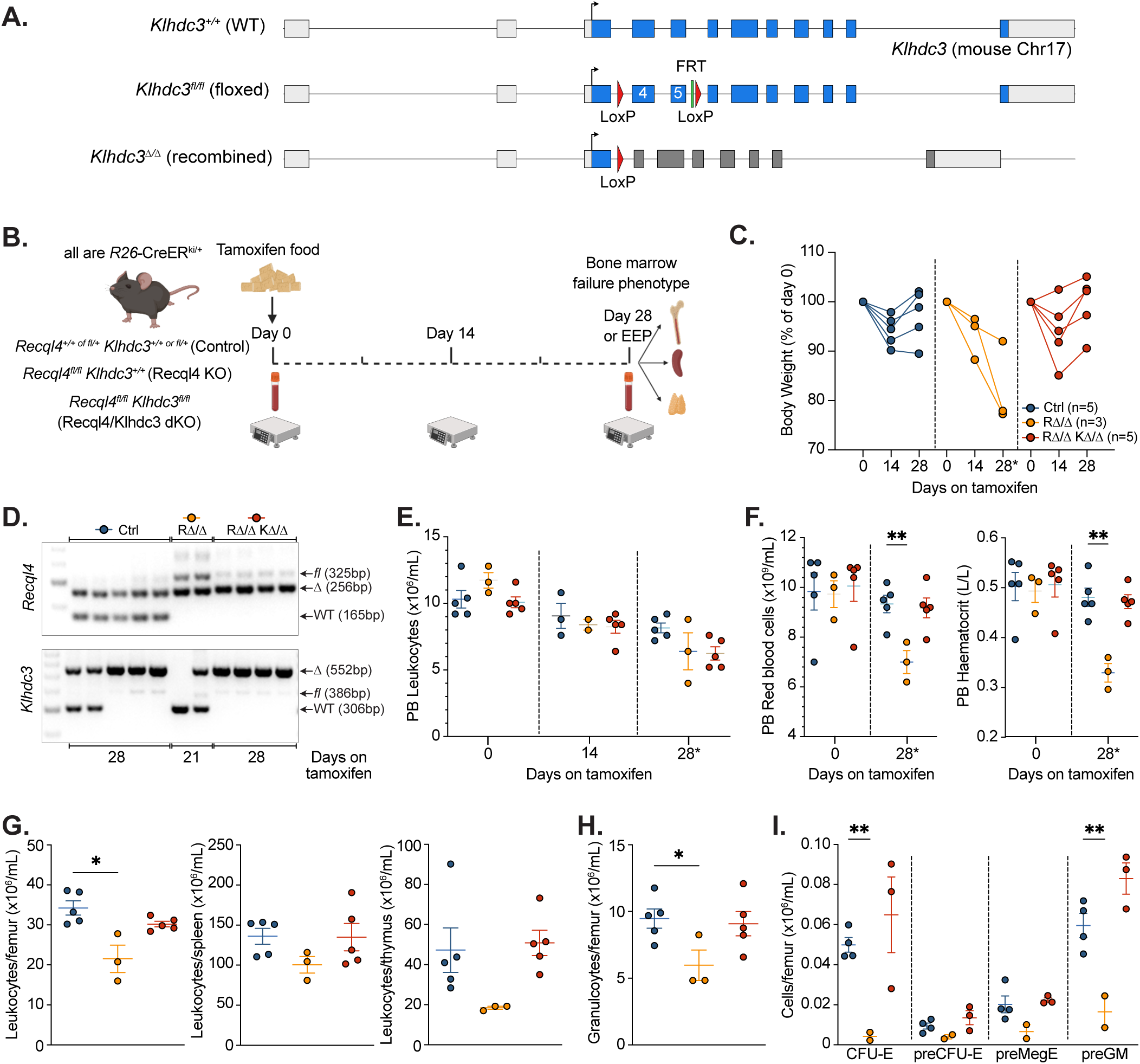
Loss of KLHDC3 prevents bone marrow failure in RECQL4 deficient mice *in vivo*. A. Schematic outline of *Klhdc3* conditional allele; exon 4 and 5 are flanked by loxP elements. B. Schematic outline of experiment. All animals are *R26*-CreER^T2Ki/+^ and were fed tamoxifen containing food *ad libitum* from Day 0. Control mice are of mixed genotypes containing wild-type or heterozygous alleles of either *Recql4* or *Klhdc3* and are littermates of the *Recql4* KO (*Recql4^fl/fl^ Klhdc3^+/+^*) and *Recql4*/*Klhdc3* dKO (*Recql4^fl/fl^ Klhdc3^fl/fl^*). Both male and female mice were used for all genotypes. All animals were assessed on day 0 and day 14 and then either at day 28 or EEP (ethical endpoint based on health assessment) which ever came first. C. Body weights of mice of each indicated genotype; each line represents an individual. D. Genotyping of genomic DNA from whole bone marrow cells for *Recql4* and *Klhdc3* recombination respectively. Days on tamoxifen indicate day of collection. E. Peripheral blood leukocyte counts at indicated day for each genotype cohort. Day 28* indicates either day 28 or EEP. F. Peripheral blood red blood cell counts and haematocrit either day 28 or EEP. G. Total leukocyte counts in per femur, spleen and thymus respectively either day 28 or EEP. H. Granulocyte numbers per femur in the bone marrow at either day 28 or EEP. I. Analysis of myelo-erythroid progenitors in the bone marrow of each genotype either day 28 or EEP. Populations assessed are within the Lineage^-^c-Kit^+^Sca-1^-^ population of the bone marrow and are the colony-forming unit erythroid (CFU-E); pre-colony-forming unit erythroid (pre-CFU-E); pre-megakaryocyte erythroid progenitor (preMegE) and pre-granulocyte macrophage progenitor (preGM). Data expressed as mean ± sem with data pooled from two independent experiments; n as indicated by each dot; *P <* 0.05, \*\**P <* 0.01 as indicated with statistical comparisons from ANOVA with multiple-comparisons correction calculated using Prism software.

Firstly, we tested whether the loss of *Klhdc3* would rescue the fully penetrant *in vivo* BM failure phenotype previously characterised in *Recql4* deficient and point mutant mice (Figure 3B) (Castillo-Tandazo *et al*., 2019; Smeets *et al*., 2014). We generated cohorts of *R26-*CreER^ki/*+*^ *Recql4^fl/fl^ Klhdc3^fl/fl^* and mixed *Recql4*/*Klhdc3* wild type and heterozygous floxed controls (littermates, co-housed and treated similarly, both males and females used), and fed them with tamoxifen containing food for up to 28 days as previously described (Figure 3B) (Castillo-Tandazo *et al*., 2019; Smeets *et al*., 2014). All mice lost weight initially, associated with the introduction of the tamoxifen diet and independent of genotype (Figure 3C). In contrast to control genotypes that fully recovered weight after the first 2 weeks of treatment, the single mutant *Recql4^1/1^* mice had a steady and progressive weight loss throughout the tamoxifen treatment period and had to be euthanised at day 21 (Ethical End Point; EEP) (Figure 3C). In accordance with our previous work, several of the *Recql4^1/1^* mice exhibited pale feet and tail tips, consistent with the development of severe anaemia that accompanies the BM failure phenotype (Castillo-Tandazo *et al*., 2019; Smeets *et al*., 2014). In contrast, the *Recql4^1/1^Klhdc3^1/1^* double mutant mice regained weight similar to control genotypes at day 28, and did not display clinical signs of anaemia. Assessment of the DNA from BM cells demonstrated that the *Recql4* alleles had complete deletion whilst *Klhdc3* was fully recombined suggesting that loss of KLHDC3 was able to protect from BM failure induced by a loss of Recql4 *in vivo* (Figure 3D). Analysis of the peripheral blood confirmed severe anaemia (reduced red blood cell numbers and haematocrit; HCT) in the single mutant *Recql4^1/1^* mice, which was prevented by concomitant loss of *Klhdc3* (Figure 3E-3F).

Similarly, total cellularity of the BM, spleen and thymus was rescued in the *Recql4^1/1^Klhdc3^1/1^* double mutant mice (Figure 3G). *Recql4^1/1^* mutant mice had reduced cellularity in the myeloid lineage and erythro-myeloid progenitors, which again was completely rescued in *Recql4^1/1^Klhdc3^1/1^* mice to comparable levels as in control mice (Figure 3H-3I).

Finally, we also looked at embryonic development. Homozygous *Recql4^-/-^*(a germ-line deleted version of the *Recql4^fl/fl^* allele used herein) is lethal before embryonic day 10.5 (E10.5) (Smeets *et al*., 2014). When *Recql4^+/-^* and *Klhdc3^+/-^* alleles were crossed, viable and macroscopically normal *Recql4^-/-^Klhdc3^-/-^*embryos were detected at the expected Mendelian ratio at E12.5, although no *Recql4^-/-^Klhdc3^-/-^* animals were identified postnatally (Figure S5) (Smeets *et al*., 2014). Collectively, these results demonstrate that KLHDC3 deletion completely prevents the development of BM failure in adult conditional RECQL4-deleted mice *in vivo* and significantly extended the survival of *Recql4^-/-^*embryos.

### KLHDC3 is a specific rescue allele and involves its E3 activity

Having established that loss of *Klhdc3* protected against the phenotypes associated with a loss or mutation of *Recql4* both in cell lines and *in vivo*, we sought to understand mechanistically how this occurred. We generated a series of KLHDC3 mutant constructs to re-express in the rescued *Recql4^1/1^Klhdc3^1/1^* mutant cell lines. We made a wild-type KLHDC3, a dominant negative (DN) point mutated KLHDC3, with mutations in the Elongin B/C box and Cullin 2 binding domain that render it unable to participate in the degradation of substrates, and a C-terminal deleted mutant than lacks the Elongin and Cullin 2 interaction domains entirely (Figure 4A) (Mahrour *et al*., 2008). The re-expression of the WT but not the mutant KLHDC3 proteins resulted in death of the *Recql4^1/1^Klhdc3^1/1^* mutant cell lines without any appreciable effect on control cells (Figure 4B). This demonstrates that the activity of KLHDC3 as a substrate recognition receptor that recognises and facilitates ubiquitin-mediated destruction of proteins with specific C-terminal degron motifs is the specific function accounting for the observed rescue.

**Figure 4.**
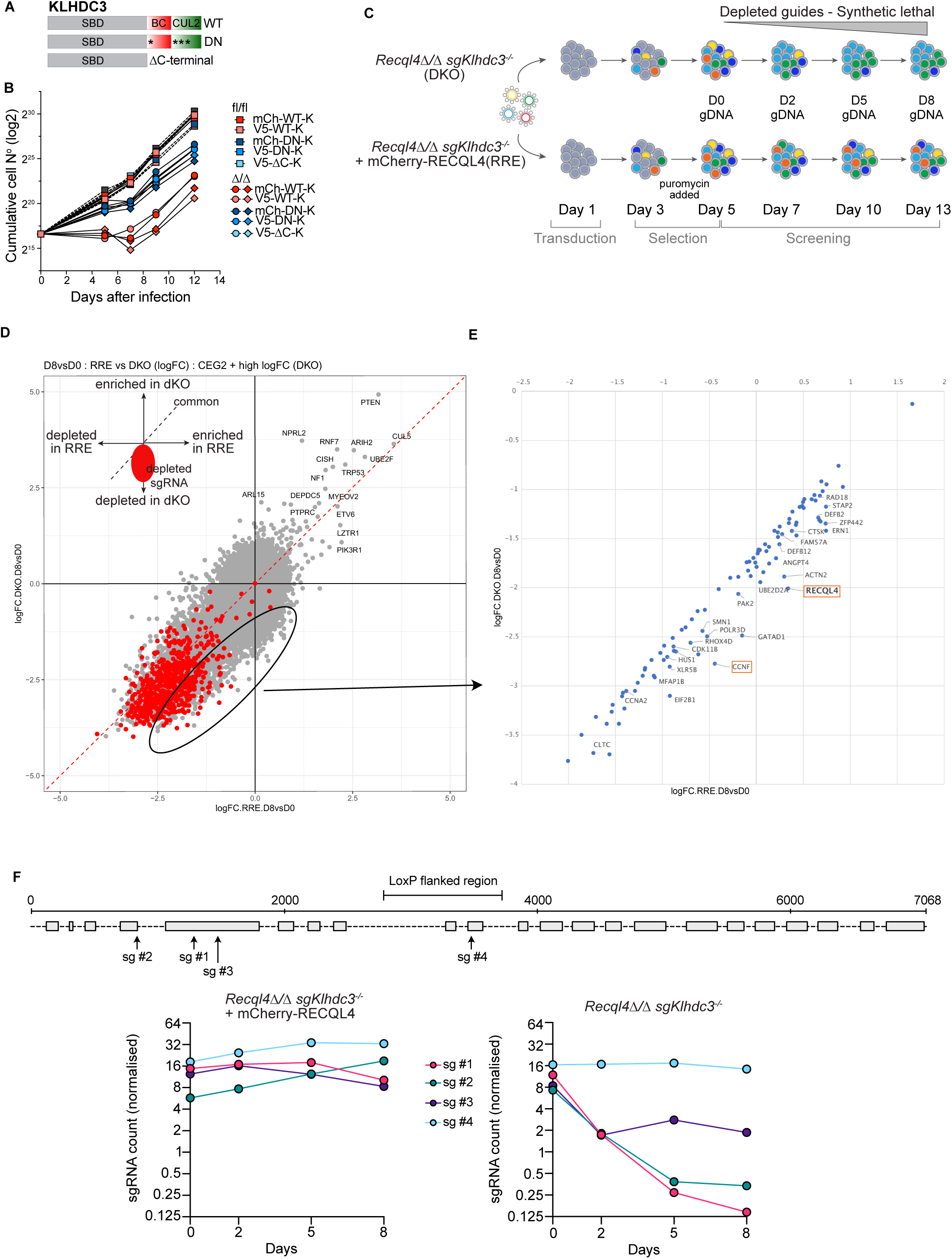
KLHDC3 is a specific rescue allele and its E3 activity is required. A. Schematic outline of KLHDC3 constructs used (WT: wild type, DN: dominant negative, *: point mutations, Δ: deletion). B. Re-expression of KLHDC3 or the DN or ΔC terminal mutant in *Recql4^fl/fl^* or *Recql4^1/1^* sg*Klhdc3* cells. Data expressed as cumulative cell number after infection with the KLHDC3 construct indicated. Data representative of three experiments using 2-4 cell lines each genotype per experiment. C. Outline of genetic screen designed to identify if loss of individual substrates of KLHDC3 would result in lethality of *Recql4^1/1^* sg*Klhdc3* cells (labelled as DKO – double knock-out) and to identify synthetic lethal interactions with *Recql4^1/1^* sg*Klhdc3*. Cells were compared to control *Recql4^1/1^* sg*Klhdc3* reconstituted with mCherry-RECQL4 (labelled as RRE – RECQL4 re-expressed). D. Summary of MaGeCK maximum likelihood estimation (mle) analysis of screen showing depletion of sgRNAs targeting the indicated genes in the *Recql4^1/1^* sg*Klhdc3* cells (DKO) compared with those reconstituted with mCherry-RECQL4 (RRE) at day 8/day 0. Red dots, high-confidence, indicate statistically significant depletion. Area of interest indicated and expanded into panel E. E. Expanded view of the sgRNAs displaying preferential depletion in the *Recql4^1/1^* sg*Klhdc3* cells. F. Schematic outline of where the individual sgRNA against *Recql4* align relative to the region deleted in the *Recql4^fl/fl^* mice (marked as LoxP flanked region). Normalised individual sgRNA counts for each indicated sgRNA against *Recql4* in the control (RRE) or *Recql4^1/1^* sg*Klhdc3* (DKO) cells. sgRNAs that are depleted target sequences 5’ to the LoxP flanked region that has been deleted in the *Recql4^1/1^* sg*Klhdc3* cells.

Having established that the rescue was specific to the substrate recognition capacity of KLHDC3, we sought to identify the key substrate(s) that were being stabilised in the absence of KLHDC3 that could explain the rescue of *Recql4^1/1^*. As computational prediction of KLHDC substrates based degron-like C-termini is notoriously unreliable (Yeh *et al*, 2021), we turned to genetic screening. We posited that deletion of candidate KLHDC3 substrate(s) in the *Recql4^1/1^Klhdc3^1/1^* mutant cell lines should revert the rescue and result in specific lethality due to the absence of RECQL4 (Figure 4C).

Cas9 expressing *Recql4^1/1^Klhdc3^1/1^* cells (referred to as DKO) and control cells complemented with WT RECQL4 (referred to as RRE; <u>R</u>ECQL4 re-<u>e</u>xpression) were infected with the Brie sgRNA library. At day 0 and days 2, 5 and 8, cells were collected, and DNA was extracted and assessed for sgRNA representation to identify sgRNA that were specifically depleted in the *Recql4^1/1^Klhdc3^1/1^* cells compared to control cells re-expressing RECQL4. This resulted in a broader range of candidate genes than the initial screen (Figure 1B; Dataset S1), including many core essential genes and those with known involvement in cell cycle control and DNA replication (Figure 4D-4E; Dataset S2).

Moreover, four of these genes, *Dna2*, *Gins4*, *Rfc3* and *Timeless*, were also recently identified in human dual-guide CRISPRi screening with RECQL4 sgRNAs in RPE-1 cells (Fielden *et al*., 2025), suggesting a conservation of genetic pathways across species (Figure S6). As an example of identified candidates, we validated synthetic lethality with loss of Cyclin F (*Ccnf*) by showing that guides targeting *Ccnf* were more rapidly depleted from the *Recql4^1/1^Klhdc3^1/1^* cells compared to control cells (Figure S7). This demonstrated that the screen was robust in resolving synthetic lethal genetic interactions with the loss of RECQL4.

Remarkably, we also identified 3 of the 4 sgRNA targeting *Recql4* amongst the most highly depleted candidates (Figure 4F). The identification of *Recql4* sgRNA as synthetic lethal interaction in the *Recql4^1/1^Klhdc3^1/1^* cells but not in the control was puzzling. The *Recql4^fl/fl^* allele used here phenocopies other reported null alleles both with respect to both germline and lineage-specific deletions (Lu *et al*., 2015; Ng *et al*., 2015). This floxed allele results in the loss of exon 9 and 10, with introduction of multiple in-frame stop codons expected to result in non-sense mediated decay of the transcript and a functionally protein null state (Castillo-Tandazo *et al*., 2019; Smeets *et al*., 2014).

Interestingly, the *Recql4* sgRNAs that demonstrated synthetic lethality targeted *Recql4* in exons 4 and 5, prior to the loxP sites that remove the protein coding region from exon 9 and 10 (Figure 4F). This was confirmed by individually targeting exons 4, 5, 10 and 15 in *Recql4^fl/fl^* control, *Klhdc3^-/-^* and *Recql4^1/1^Klhdc3^1/1^* cells. The most plausible explanation for this finding was that the residual Cre/loxP deletion product of the *Recql4^fl^* allele itself might be the target of KLHDC3, and that the stabilisation of this truncated protein product was sufficient for the observed rescue.

### KLHDC3 loss mediates rescue by stabilising the recombined RECQL4-truncation fragment

The recombined *Recql4^Δ^* mRNA transcript is predicted to retain the first 480 amino acids (aa) of RECQL4 (from exons 1-8) plus an additional frame-shifted 50aa (from exons 11 and 12), lacking the entire helicase domain and C-terminal region. Notably, the predicted C-terminal protein sequence of this 530aa protein was VPRGLGGRG*, approximating the previously characterised KLHDC3 recognition motif of RxxxxRG* (Figure 5A) (Lin *et al*, 2015; Lin *et al*., 2018; Rusnac *et al*, 2018; Timms *et al*., 2023; Yeh *et al*., 2021; Zhang *et al*., 2023).

**Figure 5.**
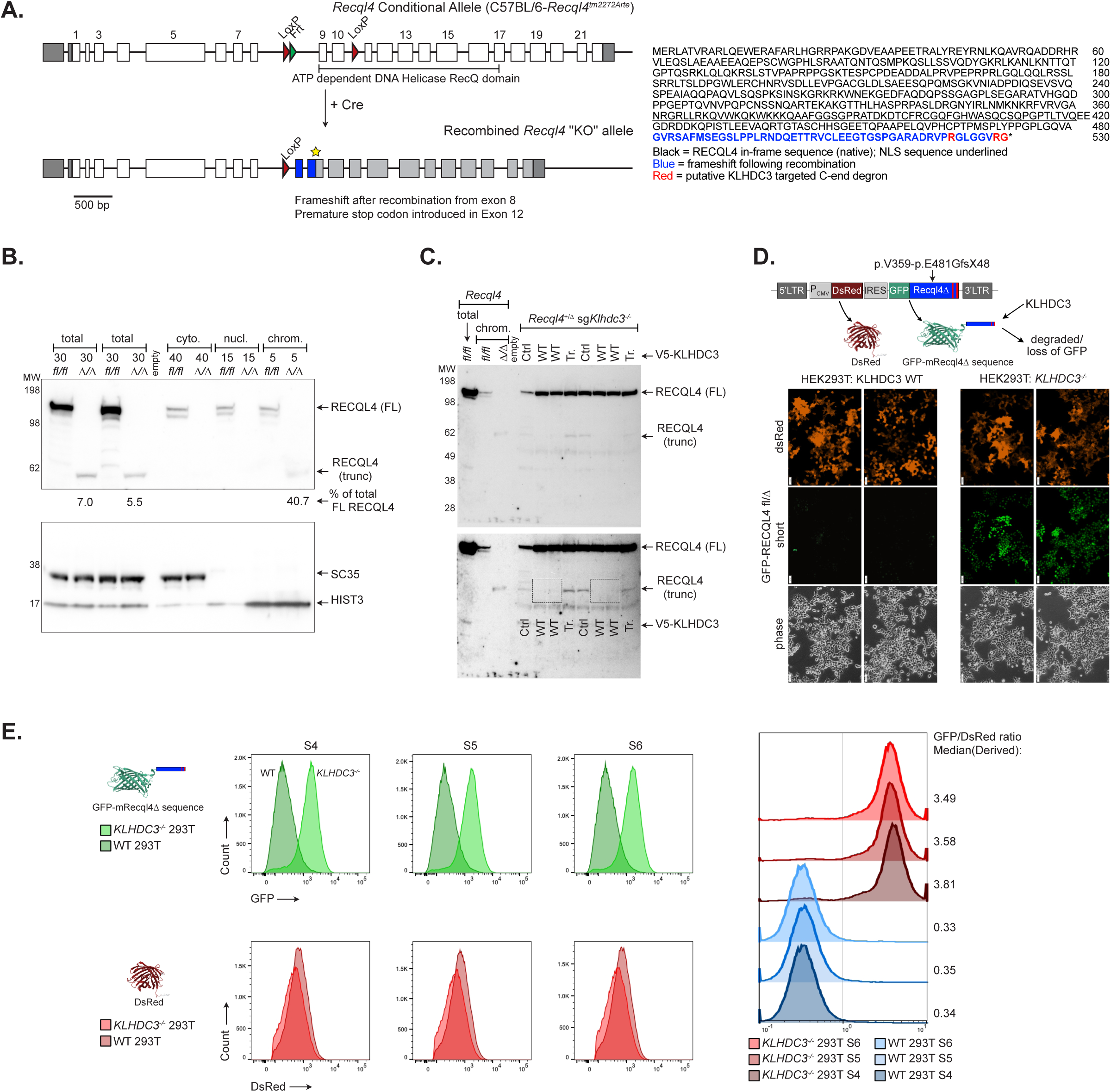
The recombined *Recql4* allele protein product is stabilised by loss of KLHDC3. A. Analysis of the predicted protein product arising from recombination of the *Recql4^fl/fl^* allele. The predicted protein product remains in frame for the first 480 amino acids (indicated in black font) then is predicted to be frameshifted (indicated in blue font) with a stop codon at 530aa. The putative KLHDC3 degron motif (RxxxxxRG*) is highlighted in red font. B. Western blot analysis of whole cell lysate or cytoplasmic, nuclear and chromatin fractions from cells of the indicated genotype assessing RECQL4 (upper panel) or SC35 or HIST3 (lower panel) expression. % of truncated RECQL4 as a percentage of full length (in fl/fl cell) as quantitated by iBright (Invitrogen/Thermo Fisher) image analysis software; fl/fl = *Recql4^fl/fl^*; Δ/Δ = *Recql4^1/1^* sg*Klhdc3.* Data representative of 2 independent replicates. C. Re-expression of wild-type full length V5-tagged KLHDC3 (either empty vector; WT1, WT2 or ΔC terminal mutant (Tr)) in *Recql4^+/-^ Klhdc3^-/-^* cells. Experiment repeated twice with the same cell line: once without MG132 (lanes 5-8) and once with MG132 (lanes 9-12). Lanes 5 and 8: Ctrl (no vector), lanes 6, 7 and 10, 11: WT KLHDC3, and lanes 8 and 12: ΔC-term KLHDC3 D. The C-terminal region of the *Recql4* deleted allele coding for p.V359-E481GfsX48 was cloned into the GPS-reporter plasmid. In this plasmid DsRed is constitutively expressed and GFP stability/expression is determined by the C-terminus of the fused RECQL4 deleted product. This reporter was expressed in KLHDC3 WT 293T cells or *KLHDC3^-/-^* 293T cells as indicated. Fluorescence signal was detected by live cell fluorescent imaging. Scale bar represents 50*μ*m. E. Flow cytometric analysis of the GFP and DsRed expression of 3 independently generated RECQL4 p.V359-E481GfsX48 clones (indicated as S4, S5 and S6 respectively) transduced in KLHDC3 WT *KLHDC3^-/-^* 293T cells. Stabilisation of GFP is quantitated by the GFP/DsRed derived median (right panel). Note the left shift in the KLHDC3 WT 293T cells indicative of destruction of the GFP-RECQL4 p.V359-E481GfsX48 GPS reporter.

Western blotting completed after the initial identification of the rescue using whole cell lysate of the *Recql4^1/1^ Klhdc3^1/1^* myeloid cells had not been able to detect a protein of the predicted size of the 530aa RECQL4 protein product of the *Recql4^Δ^* allele (Figure S2F-S2G). We revisited this with an improved sample preparation method and additional protease inhibition and were able to detect a protein of ∼58kDa, the expected mass based on the sequence (Figure 5B). This was detectable in whole cell lysates at between 5.5-7% of the levels of the endogenous full length RECQL4. As the predicted protein is expected to retain the nuclear localisation signal as part of the in-frame portion of RECQL4, we assessed where the truncated protein product was located. Based on compartmental fractionation, the truncated protein was exclusively associated with the chromatin fraction of the cell, suggesting that the *Recql4^Δ^* protein product was retained on the DNA and potentially sufficient to compensate for the absence of full length RECQL4 (Figure 5B).

Having established conditions to detect the truncated protein product, we then sought to directly establish that KLHDC3 was stabilising the 530aa protein. We re-expressed WT and C-terminal truncated mutant of KLHDC3 described earlier in *Recql4^+/1^Klhdc3^1/1^* cells (derived from bone marrow from offspring of *Recql4^+/-^*x *Klhdc3^+/-^* crosses) (Smeets *et al*., 2014). The expression of WT KLHDC3, but not C-terminal truncated, led to the loss of the detected 530aa RECQL4^Δ^ protein product (Figure 5C). These results collectively establish that the rescue of *Recql4^1/1^* cells, and point mutant cells used in the screen which also had the *Recql4^fl^* allele in them, was most likely due to KLHDC3 stabilisation of the remnant RECQL4 protein product resulting from recombination of the floxed allele.

To assess the stability of the predicted *Recql4^Δ^* protein product, we fused its last 172aa of the *Recql4^Δ^* protein product (122aa of in-frame RECQL4 and 50aa of frameshifted sequence; from position 358 of the native protein; RECQL4^Δ^ p.358-530aa) to an N-terminal GFP, and compared its stability to the constitutively expressed DsRed using the global protein stability (GPS) assay (Figure 5D) (Yen & Elledge, 2008; Yen *et al*., 2008). The ratio of GFP to DsRed reflects of the stability of the C-end degron fused to GFP. When expressed in KLHDC3 wild-type HEK293T cells we could detect robust DsRed signal but no GFP by either microscopy (Figure 5D) or flow cytometry (Figure 5E). We then expressed the construct in *KLHDC3^-/-^* HEK293T cells (Figure S8A) and obtained robust expression of GFP (Figure 5D-5E; Figure S8A). We confirmed the specificity of *KLHDC3^-/-^*HEK293T cells using previously characterised GPS-reporter constructs that are targeted by KLHDC3 or APPBP2 respectively (Figure S8B-S8C) (Huang *et al*., 2023; Lin *et al*., 2018; Yeh *et al*., 2021). This formally established that the transcript arising from the *Recql4^Δ^* allele, if translated, was forming a C-end degron recognised by KLHDC3 and specifically stabilised by loss of KLHDC3. These results collectively demonstrate that the rescue of *Recql4^1/1^* and point mutant cells, which also contain the *Recql4^fl^* allele, was due to KLHDC3 deletion-induced stabilisation of the remnant RECQL4 protein product resulting from recombination of the floxed allele.

### Genome-wide replication timing is maintained in the absence of the RECQL4 ATP Helicase domain and activity

A recent study suggested that, unlike what was previously reported for RECQL4 p.K525A ATP-binding mutant mice (Castillo-Tandazo *et al*., 2019), there was a requirement for the ATP-dependent helicase activity of xRECQL4 to support normal DNA replication in metazoans (Terui *et al*., 2024). This study used single molecule resolution analysis in *Xenopus* oocyte extracts to conclude that the ATP-dependent activity of RECQL4 was required to efficiently remove DONSON from the Cdc45/Mcm2-7/GINS (CMG) helicase to allow DNA replication to proceed. The KLHDC3 loss mediated rescue of the myeloid cell lines allowed us to directly address this question, as the 530aa RECQL4^Δ^ protein product does not encode the ATP-dependent helicase region nor the entire C-terminal domains of RECQL4. Therefore, if the ATP-dependent helicase activity was essential, we should see significant changes in replication timing in the *Recql4^1/1^Klhdc3^1/1^* rescued cells. We directly assessed genome-wide DNA replication kinetics using whole genome sequencing and the Timing Inferred from Genome Replication (TIGER) computational analysis method in WT and *Recql4^1/1^Klhdc3^1/1^* cells (Koren *et al*., 2021). We resolved high resolution (10kb) maps of DNA replication timing in all cell lines assessed and from this were able to compare replication timing in the presence of full length RECQL4 (WT control cells) or when there was no ATP-dependent helicase region at all (*Recql4^1/1^Klhdc3^1/1^* cells) (Figure 6A-6D; Figure S9) (Koren *et al*., 2021). This analysis demonstrated that there were, at most, subtle differences between cells with and without the ATP-dependent helicase and C-terminal domains of RECQL4 (Figure 6D). When assessed across individual chromosomes there was only a small variation apparent for the majority of chromosomes (Figure S9). There was more variability in the *Recql4^1/1^Klhdc3^1/1^* cells compared to the controls, potentially derived from sub-clonal variance that is apparent in the compound mutants (Figure 6D, Figure S9). This analysis, whilst not able to completely exclude true difference in replication timing, indicate that any difference at most is small.

**Figure 6.**
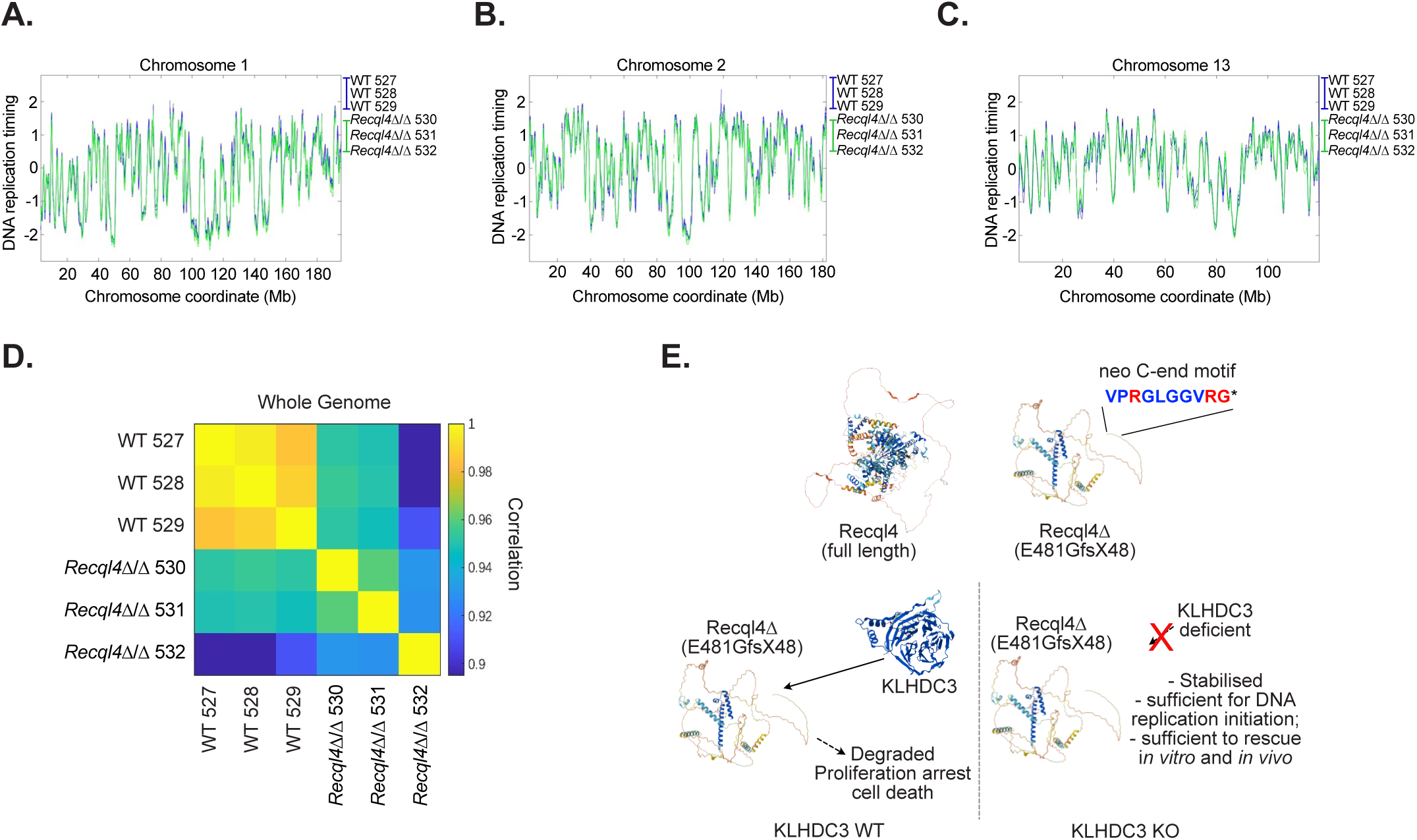
Normal genome-wide DNA replication timing does not require the helicase domain of RECQL4. A. Computed DNA replication timing across chromosome 1 with each genotype averaged to a single line per genotype (blue = WT; green = *Recql4^1/1^* sg*Klhdc3* cells). B. Calculated DNA replication timing for chromosome 2. C. Calculated DNA replication timing for chromosome 13. D. Correlation analysis of DNA replication timing across the whole genome. E. Graphical summary of the results. In wild-type cells, the protein product from the *Recql4^fl/fl^*allele recombination (*Recql4^1/1^;* RECQL4 p.E481GfsX48) is unstable and subjected to targeting by KLHDC3 due to the formation of a specific neo C-end degron motif recognised by KLHDC3. This leads to ubiquitin mediated proteasomal degradation of this RECQL4 p.E481GfsX48 protein and failure of DNA replication. When KLHDC3 is deleted, the RECQL4 p.E481GfsX48 protein is stabilised and retained on chromatin where it is sufficient to enable near normal DNA replication and cell division.

Therefore, we can conclude that the ATP-dependent helicase and C-terminal domains of RECQL4 are not essential for DNA replication to proceed in an ordered and efficient manner in the myeloid cell lines. In addition, the complete protection from BM failure in the *Recql4^1/1^Klhdc3^1/1^* animals further suggests that this also applies *in vivo*, consistent with our previous *in vivo* work demonstrating that the ATP-dependent helicase function of RECQL4 was not essential in mammals (Castillo-Tandazo *et al*., 2019).

## Discussion

Mutations in RECQL4 are associated with Rothmund-Thomson Syndrome (RTS) type II, a syndrome characterised by both relatively benign pathologies, including low bone mass and immune compromise, and a highly elevated rate of cancer, particularly osteosarcoma and a range of other malignancies including haematological cancers (Larizza *et al*, 2010; Wang *et al*., 2001). Despite this association with severe human disease, the fundamental biology of RECQL4 and our understanding of its genetic interactions is very restricted compared to the related RecQ helicases such as BLM. RECQL4 was thought to be the mammalian homologue of the essential yeast protein *Sld2*. However, the mammalian *Sld2* homologue has recently been identified as DONSON (Cvetkovic *et al*., 2023; Evrin *et al*., 2023; Hashimoto *et al*., 2023; Kingsley *et al*., 2023; Lim *et al*., 2023). Therefore, significant gaps remain in our understanding of RECQL4’s function. We sought to address this knowledge gap using forward genetic screens and human like mutations engineered in mouse cells and *in vivo* in murine models. Whilst our study yielded an unexpected finding specifically related to models we used, it has allowed us to make significant advances in understanding both the genetic interactions and functions of RECQL4 protein in mammals.

We undertook a genome-wide suppressor screen to identify factors whose loss restored proliferative capacity and viability to cells with a non-viable RECQL4 truncating mutation (Castillo-Tandazo *et al*., 2021; Castillo-Tandazo *et al*., 2019). This resulted in the identification of loss of KLHDC3 as a strong suppressor, including for *in vivo* phenotypes. KLHDC3 is a substrate recognition component of the CRL E3 ubiquitination system (Pilcher *et al*., 2025; Rusnac *et al*., 2018; Timms *et al*., 2023; Yeh *et al*., 2021; Zhang *et al*., 2023). KLHDC3 targets proteins that have RxxxG C-end degron motifs, promoting their ubiquitination and degradation (Timms *et al*., 2023; Yeh *et al*., 2021; Zhang *et al*., 2023). Further confirming the specificity of the KLHDC3 enrichment, we failed to observe any significant enrichment of sgRNA against other closely related KLHDC family members, such as KLHDC2 or KLHDC10, or against the scaffold components that the KLHDC proteins interact with (e.g. Elongin B/D, Cullin 2). However, this rescue was specific to the design of the floxed *Recql4* allele in our mice (Figure 6E) (Smeets *et al*., 2014). Although this limited the generalisability and involvement of KHLDC3 in the fundamental biological roles of RECQL4, and by extension in the pathology of RTS, our finding that minute amounts of a severely truncated RECQL4 protein fragment largely supports its essential role in DNA replication allows for important general conclusions.

The data indicate that a significantly truncated RECQL4 protein, only retaining the first in-frame 480aa, was present in the rescued cells at less than 10% of the levels of WT. The truncated protein was found, near exclusively, in the chromatin fraction of the nucleus suggesting that it remains predominantly associated with chromatin at comparable molar levels to chromatin-bound wildtype full-length RECQL4. The capacity of this residual level of RECQL4 to be functionally relevant is consistent with the results obtained in *Drosophila* where mutants that retained less than 7.0% of RECQL4 protein remained largely normal (Wu *et al*, 2008). This suggests that only a small fraction of the endogenous level of RECQL4 is required to support DNA replication and homeostasis. This has important implications for studies using siRNA and other efficient, but not absolute, means to reduce expression of *Recql4* (Padayachy *et al*., 2024; Terui *et al*., 2024). This is also the case for experimental systems that utilise depletion, where even trace amounts of functionally active RECQL4 may confound interpretation of results. The predicted protein produced from the deleted *Recql4* allele retains the N-terminal in-frame up to aa 480, including regions that would contain the phosphorylation sites recently identified in RECQL4 to regulate baseline and dormant replication origins (Thakur *et al*., 2025). We could show that homozygous RECQL4 p.E481GfsX48 (the protein product from the recombined *Recql4* floxed allele) was sufficient for normal DNA replication and prevented BM failure *in vivo* if stabilised by the loss of KLHDC3. Our previous results showed that RECQL4 p.G522EfsX43 and RECQL4 p.R347X variants did not rescue this *in vivo* phenotype (Castillo-Tandazo *et al*., 2021; Castillo-Tandazo *et al*., 2019). The G522Efs protein product, longer in predicted region of retained RECQL4 in-frame sequence, is not stable and the R347X protein, although stable and localised in the nucleus, lacks the Zn-knuckle motif and most of its upstream region previously shown to be required for ssDNA and fork DNA substrate binding (Marino *et al*, 2016). In contrast, retention of a minute level of stable protein expression encoding the first 480aa of RECQL4 was able to substitute for full length RECQL4 in many settings.

The mechanism via which the RECQL4 p.E481GfsX48 enables relatively normal DNA replication is still to be fully resolved. Our analysis indicate that this small fragment supports complete DNA replication, but delays completion of DNA synthesis with a shift towards late S phase and even into M phase. This shifted DNA synthesis, evidenced by EdU, presumably depends on MiDAS contributing to the ability of the cells to complete replication (Bhowmick *et al*., 2023). This may occur in a stochastic manner within a population of *Recql4^1/1^Klhdc3^1/1^* cells leading to genomic alterations, which may be why we do not observe a global replication timing difference in bulk analysis. If this also occurred stochastically across the genome, i.e., different locations in every cell and not necessarily during every cell cycle, then that would explain why we see no replication timing variation in bulk analysis, and could further explain why we nonetheless do see a replication phenotype in as measured by EdU/Ki67 and MCM3 chromatin loading. Recent work demonstrated that RECQL4 was required for replication origin choice, and the choice between dormant vs. baseline origins, and the replication stress response (Thakur *et al*., 2025). The small RECQL4 fragment is associated with an increase in licensed replication origins. This may be because the helicase domain normally acts to limit how many origins are licensed (Terui *et al*., 2024). As a consequence, more active origins during early S phase with increased dNTP depletion may cause the replication stress that ultimately delays complete replication into M phase. Alternatively, increased origin licensing could also be a compensatory response where more active origins result in shorter replicons making up for slower replication in the absence of the helicase domain.

Remarkably, the low levels of RECQL4 p.E481GfsX48 product of the *Recql4^Δ^* allele in the absence of KLHDC3 were fully sufficient to support adult haematopoiesis and prevent BM failure resulting from loss of *Recql4 in vivo* (Smeets *et al*., 2014). In contrast, while it significantly extends the embryonic viability of *Recql4*-mutated embryos, it does not support full completion of embryonic development. This difference in the extent of *in vivo* rescue may reflect the need for highly proportional and synchronised cell proliferation during embryonic development that may be less tolerant of altered, even subtly, replication timing than the more asynchronous process of haematopoiesis. The unanticipated generation of a highly specific C-terminal degron sequence as favoured by KLHDC3 could not have been predicted when the *Recql4^fl^* allele was made, as KLHDC3 and the C-end degron model had not been described at the time (Koren *et al*., 2018; Lin *et al*., 2018; Smeets *et al*., 2014; Timms & Koren, 2020). This would mean that in the *Recql4^fl/R347X^* cells used for the original screen, the RECQL4 p.E481GfsX48 product of the *Recql4^Δ^* allele was responsible for mediating the rescue even in the presence of the p.R347X protein product. Our findings emphasise that the nature and basis of genetic interactions observed in genome-wide screens, both *in vitro* as here but increasingly *in vivo*, needs to be carefully validated and considered to demonstrate that it is a specific finding related to biological function, not one related to the specific model or genetic modifications.

Another key finding of this study is that using a genome-wide screening we were unable to identify any additional loss of function alleles, aside from KLHDC3 and the mutant RECQL4 protein product mechanism described, that could rescue the proliferation and viability defects of cells expressing a RECQL4 mutant protein similar to those reported in patients with RTS. This result indicates that there are unlikely to be monogenic loss of function rescue alleles for RECQL4 mutation, suggesting that RECQL4 is an essential and non-redundant regulator of DNA replication and viability. Therefore, it is improbable that there will be genetic means to bypass the molecular consequences that occur when RECQL4 is mutated as in RTS patients. This has significant implications for how we consider future therapy for RTS patients with RECQL4 mutations and indicates that direct correction of the mutations in RECQL4 is likely the most feasible path to treating or curing these patients.

Heterozygosity for RECQL4 mutation is well tolerated, based on the absence of phenotypes in the parents of RTS patients, suggesting correction of only one allele of RECQL4 is likely to be significantly beneficial. The rapid advances in the potential of genetic corrective therapy, such as CRISPR or redirected RNA editing, offer potential pathways to explore. Collectively the present results, together with our previous work, do not support an essential function for the ATP-dependent helicase activity of RECQL4 in mammalian DNA replication.

## Acknowledgements

The authors thank L Purton, JX Xu and N Hoch for comments and discussion; K Simpson, T Gulati and H Beetham (Victorian Centre for Functional Genomics, Peter MacCallum Cancer Centre, Australia) for providing Brie sgRNA lentiviral library aliquots; WEHI Antibody Facility and Monash Antibody Technology Facility (MATF), Monash University for purification of RECQL4 antibody; BGI and Novogene for sequencing services; H-C S Yen (Academia Sinica Institute of Molecular Biology, Taipei) for GPS reporter plasmids; M Kamps (UCSD, USA) for HoxB8 plasmid; Addgene for plasmid distribution as noted; St. Vincent’s Hospital Bioresource’s Centre for care of experimental animals; S Taylor, A Goradia and E Tonkin for technical assistance.

The authors acknowledge the facilities, and the scientific and technical assistance of the following Phenomics Australia nodes: Victorian Centre for Functional Genomics, Peter MacCallum Cancer Centre; and the Rodent Histopathology Service, University of Melbourne. The *Klhdc3^fl/fl^* mice were produced via CRISPR genome editing by the Monash Genome Modification Platform (MGMP), Monash University is a node of Phenomics Australia. Schematic figures were made BioRender.com and NIH Bioart.

## Author contributions

Conceptualization: MFS, TYT, CRW

Methodology: PAB, WJC-T, AMC, JH, AK, MFS, CRW

Investigation: PAB, WJC-T, AMC, CP, JKH, JH, AK, MFS, CRW

Visualization: PAB, WJC-T, AMC, CP, JH, AK, MFS, CRW

Funding acquisition: JKH, TYT, CRW

Supervision: JKH, MFS, CRW Writing – original draft: MFS, CRW

Writing – review & editing: all authors

## Materials and Methods

### Ethics Statement

All animal experiments were performed according to the National Health and Medical Research Council Act 1992 and the Australian Code for the Care and Use of Animals for Scientific Purposes (2013). The procedures were approved by the Animal Ethics Committee, St. Vincent’s Hospital, Melbourne, Australia (#011/15, and 015/17). Animals were euthanased by CO_2_ asphyxiation or cervical dislocation.

### Mice

All animals were housed at the Bioresources Centre (BRC) located at St. Vincent’s Hospital, Melbourne. Mice were maintained and bred in microisolators under specific pathogen-free conditions with autoclaved food and acidified water provided *ad libitum*. The *Recql4^R347X^ and Recql4^G522Efs^* mutations and *Recql4^fl/fl^* mice (C57BL/6-*Recql4^tm2272Arte^*) have been all been previously described (Castillo-Tandazo *et al*, 2021; Castillo-Tandazo *et al*., 2019; Ng *et al*, 2015; Smeets *et al*., 2014).

*Rosa26*-CreER^T2^ mice on a C57Bl/6 background were purchased from The Jackson Laboratory (B6.129-Gt(ROSA)26Sor^tm1(cre/ERT2)Tyj^/J, Stock Number: 008463) and have been previously described(Smeets *et al*., 2014). All genotyping was performed as previously described (Castillo-Tandazo *et al*., 2021; Castillo-Tandazo *et al*., 2019; Ng *et al*., 2015; Smeets *et al*., 2014). ENU mutants were outcrossed at least 6 times and assessed across multiple generations to eliminate effects of any additional mutations. All lines were on a backcrossed C57Bl/6 background.

*Klhdc3* conditional (*Klhdc3^fl/fl^*) mice were generated by the Monash Genome Modification platform by CRISPR/Cas9 mediated insertion of loxP elements flanking Klhdc3 exons 4 and 5. A ssDNA repair template was prepared for the required sequence to insert. Cas9 protein (80ng/*μ*l; Integrated DNA Technologies (IDT)) was incubated with sgRNA (80ng/*μ*l; crRNA and tracrRNA; IDTDna) to generate RNP. The RNP and the ssDNA repair template (50ng/ml) were microinjected into C57BL/6J zygotes at the pronuclei stage. Microinjected zygotes were transferred into the uterus of pseudo pregnant F1 females. Correct insertion was screened by genomic DNA PCR and then digital droplet PCR (Bio-Rad). Correctly targeted males were identified and bred to C57BL/6J females to ensure germ-line transmission. During screening of progeny, a male with a deletion encompassing the loxP flanked region was identified (referred to as *Klhdc3^+/-^*). The male was bred to C57BL/6J females to ensure germ-line transmission. All lines were confirmed to have the expected mutations/inserted elements by Sanger sequencing of the loxP elements and the deletion event, respectively. Genotyping is described in the supplemental information and a full description of the *Klhdc3* targeted allele and phenotype is provided in a separate report.

Where indicated, tamoxifen containing diet was prepared by Specialty Feeds (Perth, Australia) containing 400mg/kg tamoxifen citrate (Selleckchem) in a base of standard mouse chow (irradiated)(Smeets *et al*., 2014). When used, tamoxifen containing chow was fed *ad libitum* for the duration of treatment.

### Generation of retrovirus/lentivirus

Retrovirus or lentivirus was produced using transient transfection of HEK293T cells as previously described (Heraud-Farlow *et al*, 2024; Smeets *et al*., 2014; Xu *et al*, 2022). Virus containing supernatant was collected at 48 and 72 hours in 1.0mL aliquots and stored at-80°C.

### Generation of Hoxb8 immortalised and Cas9 expressing cell lines

*Hoxb8 immortalised cell lines* – Bone marrow cells were collected from *R26*-CreER^T2^ *Recql4^fl/+^* (control, became *Recql4* heterozygous after tamoxifen treatment), *R26*-CreER^T2^ *Recql4^fl/fl^* (*Recql4* deficient) and *R26*-CreER^T2^ *Recql4^fl/R347X^* (expressed only the *Recql4* R347X allele after tamoxifen treatment), Ficoll density gradient separated and cultured for 48 hours in complete IMDM (20% FBS, 1% Penicillin/Streptomycin, 1% glutamine) containing recombinant mouse stem cell factor (rmSCF, 50ng/mL, Peprotech), recombinant mouse interleukin 3 (rmIL-3, 10ng/mL, Peprotech) and recombinant human interleukin 6 (rhIL-6, 10ng/mL, Amgen). After 48 hours culture, 1×10^6^ cells were spin-infected with Hoxb8 retrovirus (Wang *et al*, 2006; Xu *et al*., 2022) (Hoxb8 plasmids generously provided by Mark Kamps, UCSD) and polybrene at 1,100g for 90 minutes. After 48 hours, the non-adherent cells were passaged into complete IMDM (10% FBS, 1% Penicillin/Streptomycin, 1% glutamine) with rmSCF, rmIL-3, rhIL-6 and 1% granulocyte-macrophage colony-stimulating factor (GM-CSF) conditioned medium (from BHK-HM5 cell conditioned medium) (Xu *et al*., 2022). After 1 to 2 passages, cells were cultured in complete IMDM with only 1% GM-CSF conditioned media. The cells were fully immortalised after four weeks and were GM-CSF dependent.

*Cas9 expressing cell lines* – Following establishment, Hoxb8 immortalised cell lines were infected with Cas9 expressing lentivirus (p.lentiCas9-Blast was a gift from Feng Zhang; Addgene plasmid # 52962) (Sanjana *et al*, 2014) by spin-infection at 1,100g for 90 minutes. The infected cells were then cultured in media (IMDM, 10%FBS, 1% GM-CSF and 3µg/mL blasticidin) for two weeks to select for a Cas9-expressing population. To assess the target editing efficiency of the Cas9 Hoxb8 *R26*CreER^T2^ *Recql4^fl/+^* and *R26*CreER^T2^ *Recql4^fl/R347X^* cell lines, stable Cas9 cells were transduced with pLentiGuide (VCFG, from Addgene) expressing sgRNA targeting CD44, a cell surface marker highly expressed on all immortalised Hoxb8 cells. Forty-eight hours after infection, puromycin (0.25µg/mL) was added to select for pLentiGuide sg*Cd44* expression. Four days later, 0.5-1×10^6^ cells were stained with a PE-Cy7 conjugated anti-CD44 antibody (1:400) and analysed by flow cytometry using the FACS BD LSR II Fortessa system.

Oligonucleotide sequences (sgRNAs) and genotyping primer sequences are provided in the Supplemental information.

### CRISPR knockout pooled library screen

For loss of function rescue screening, 80×10^6^ Cas9 Hoxb8 *R26*CreER^T2^ *Recql4^fl/+^* and *R26*CreER^T2^ *Recql4^fl/R347X^*cell lines were infected with the Brie CRISPR knockout pooled library (Mouse Brie CRISPR knockout in lentiGuide-Puro pooled library was a gift from David Root and John Doench (Addgene #73633; http://n2t.net/addgene:73633; RRID:Addgene_73633) (Doench *et al*, 2016). Brie CRISPR knockout pooled library virus was generated by the Victorian Centre for Functional Genomics (RRID:SCR_025582) and provided as aliquots (Heraud-Farlow *et al*., 2024; Xu *et al*., 2022). The number of cells infected was calculated by multiplying the number of guides in the library (78,637) by the desired number of cells for each guide at the start of the experiment (aiming for 400 copies per sgRNA), divided by the multiplicity of infection required (MOI=0.3). The infection mix was centrifuged at 1,100g and 25°C for 1.5 hours and then returned to the 37°C incubator. At the end of the day, each flask was topped up with 30mL of pre-warmed medium (IMDM, 10%FBS, 1% GM-CSF and 3µg/mL blasticidin). On day 3, cells were counted and expanded to T175 flasks containing 125mL of medium at a concentration of 1×10^6^ cells/mL. Puromycin was added at a dose of 0.25µg/mL. On day 4, cells were further diluted to reach a concentration of 3.3×10^5^ cells/mL of medium plus puromycin. On day 7, puromycin was removed from the medium and cells were re-seeded at 2×10^5^ cells/mL in tamoxifen containing medium (400nM/mL) to induce CreER-mediated recombination (Day 0 designated the day that tamoxifen was added). During the next 14 days, cells were passaged every two or three days at a density to maintain at least 400 cells per sgRNA in the presence of tamoxifen, which corresponded to 2 x T175 flasks per cell line with 250mL total at a concentration of 1 or 2×10^5^ cells/mL. Additionally, at each indicated timepoint (Day 0, 4 and 9 post tamoxifen addition), pellets of 2 and 25 million cells were kept for DNA recombination analysis and gDNA isolation for library sequencing, respectively (Heraud-Farlow *et al*., 2024; Xu *et al*., 2022).

For synthetic lethal screening (Xu *et al*., 2022), the experimental procedure was similar to that described above for loss of function rescue screening with the following modifications. The Brie CRISPR knockout sgRNA pooled library virus used. The cell lines used for synthetic lethal screening were Cas9 Hoxb8 *R26*CreER^T2^ *Recql4^Δ/Δ^* sg*Klhdc3* clone 185K1B12 (see generation of clonal cell lines) and clone 185K1B12 complemented with mCherry-*Recql4*. 130×10^6^ cells per cell line were infected with the Brie CRISPR sgRNA library at an MOI of 0.3 to have 500 cells per sgRNA and cells were maintained at ≥500 cells per sgRNA by passaging 45×10^6^ cells at 2 or ∼0.8 x10^5^ cells/mL every two or three days. At least 2 pellets of 25×10^6^ cells each cell line were collected on days 0, 2, 5 and 8 (Day 0 being the third day after puromycin selection) for gDNA to assess sgRNA copy number.

### CRIPSR screen sample preparation and sgRNA library sequencing

Genomic DNA was extracted from the cell pellets using the Gentra Puregene kit (Qiagen). DNA was quantified on a Nanodrop spectrophotometer. DNA libraries were generated by PCR amplification of the integrated sgRNA constructs (as described in https://portals.broadinstitute.org/gpp/public/resources/protocols). The resultant libraries were generated and sequenced on the Illumina platform using 150bp paired end reads by Novogene (Singapore).

CRISPR screen analysis was performed using the CRISPRBetaBinomial package (CB^2^) (v 1.3.0) (Heraud-Farlow *et al*., 2024; Jeong *et al*, 2019; Li *et al*, 2015; Xu *et al*., 2022). Briefly, raw reads were mapped to the broadgpp_brie_crispr library with run_sgrna_quant function and read counts were normalised with the get_CPM function. Gene level statistics were calculated using the measure_gene_stats function. Gene level statistics were calculated using measure_gene_stats function. Annotation with ENSEMBL was performed with biomaRt (Durinck *et al*, 2009a; Durinck *et al*, 2009b). Plots were generated using ggplot2 (Wickham, 2016). Core essential genes version 2 (CEG2) was obtained from CEG2 supplemental table 2 “List of the human core essential gene set version 2” (Hart *et al*, 2017).

### Loss of function screen validation

For loss of function rescue validation, four individual sgRNAs targeting *Klhdc3* were selected and cloned into the pLentiGuide, pLentiCrispr puromycin or pLentiCrispr hygromycin vectors (sgRNA from IDTDna). mCherry-tagged codon optimised mouse *Recql4* cDNA (Castillo-Tandazo *et al*., 2021) was cloned into pLenti SFFV SV40 blasticidin or zeocin vectors (Twist Bioscience). Plasmid DNA was purified using the Isolate II Plasmid Mini kit (Bioline) and quantified using the Nanodrop spectrophotometer system for use in viral production and subsequent cell proliferation assays. The sgRNA plasmids targeting *Klhdc3* and control sgRNAs were individually transfected into HEK293T cells and transduced in Cas9 Hoxb8 *R26*-CreER^T2^ *Recql4^fl/R347X^* cells (Re-validated with two guides in fl/+, fl/R and fl/fl). Three days after transduction, puromycin was added at a dose of 0.25µg/mL for four days for positive selection. After generating cells expressing lentivirus containing sgRNAs targeting the *Klhdc3* gene, cells received tamoxifen (400nM/mL) for 14 days in a cell proliferation assay. Every two to three days, cells were counted, and the corresponding volume for the number of cells needed to re-seed 1-2×10^5^ cells/mL were centrifuged, the media discarded, and the cell pellet was resuspended in 5mL fresh tamoxifen containing medium. Cells were treated for 14 days, and at each time point, cell pellets were collected for DNA recombination/*Recql4* deletion analysis. As a control, untreated cells were kept in parallel and seeded at the same concentration.

CRISPR-Cas9 cutting and KO efficiency was determined following PCR amplification of the targeted genomic region by T7 Endonuclease I assay (T7E1, NEB M0302) according to the NEB protocol and by Sanger sequencing plus TIDE analysis (tide.nki.nl)(Brinkman *et al*, 2018). Reduced expression levels were confirmed by RT-qPCR. All primers are described in the supplemental information.

For synthetic lethal validation, three guides targeting *Ccnf* (Table S1) were cloned into pLentiCrispr puromycin vectors and transduced in Cas9 Hoxb8 *R26*-CreER^T2^ *Recql4^fl/fl^* control cells and *Recql4^Δ/Δ^* sg*Klhdc3* clones 185K1B12 (-/+ mCherry-*Recql4*) and 186K1E1 (see below). Two days after infection, puromycin was added to the cells for three days to eliminate non-infected cells. From then on cells were counted and passaged every two to three days for 11 days, and cell pellets were collected for gDNA extraction and assessment of CRISPR-induced knockout frequency by TIDE (Brinkman *et al*., 2018) and ICE (accessed via Synthego)(Conant *et al*, 2022).

### Generation of clonal cell lines

Immortalised mouse myeloid *Recql4^Δ/Δ^* sg*Klhdc3* clonal cell lines were generated at the end of a 14-day tamoxifen treatment as described in Heraud-Farlow et al(Heraud-Farlow *et al*., 2024). After Sanger sequencing, mutant clones containing homozygous indels predicted to result in loss of function, were selected for further experiments. Specifically, clone 185K1B12 containing a 38bp deletion in exon 7 predicted to result in KLHDC3 p.I216LfsX7 and clone 186K1E1, with a 391bp deletion in intron 5-exon 7, resulting in KLHDC3 p.A150X.

Human HEK293T cells transduced with pLentiCrispr sg*KLHDC3* were single-cell cloned by limiting dilution in 96-well multi-well plates after 14 days of hygromycin selection, then expanded and analysed as above. Clones 293T K1E3, K2D8 and K2D10 are described in Figure S8.

### Mouse KLHDC3 re-expression experiments

Mouse codon-optimised dominant negative (DN) *Klhdc3* was obtained from Twist Bioscience as a clonal gene in pLenti SFFV IRES puromycin with an N-terminal V5-APEX2 fusion, a L338A mutation in the BC box and S354A, C355A, L356A and P357A mutations in the CUL2 box. These mutations have been shown to abolish Elongin B/C binding in other BC box proteins or CUL2 binding in KLHDC3 specifically (Mahrour *et al*, 2008). Twist gene fragments were subsequently used to create a codon-optimised wild-type (WT) and a C-terminally truncated (p.S335X) *Klhdc3*, lacking the BC box, CUL2 box and last 17aa. These plasmids were then used to construct V5-only and mCherry-tagged versions by standard cloning techniques.

For proliferation assays, two parental *Recql4^fl/fl^* cell lines (185 and 186) and *Recql4^Δ/Δ^* sg*Klhdc3* clones 185K1B12 and 186K1E1 were transduced with V5-or mCherry tagged WT, DN and ΔC-terminal KLHDC3 expressing constructs. Puromycin was added two days after infection for three days and cells were subsequently passaged and counted every two to three days to assess survival and proliferation. For western blot analysis *Recql4^+/Δ^ Klhdc3^-/-^* cells were used to keep the cells alive after re-expression of functional KLHDC3 by one copy of WT *Recql4*. Cells were selected with puromycin two days after transduction and collected for protein as described below.

### Protein extraction and western blotting

Cell lysates were prepared in RIPA buffer (50mM Tris, 150mM NaCl, 1% NP-40, 0.5% sodium deoxycholate, 0.1% SDS, pH8.0) plus 1x Halt^TM^ (Thermo Fisher Scientific, 78440) protease and phosphatase inhibitors. Myeloid cells were pre-treated with 4-(2-aminoethyl)

Benzenesulfonyl fluoride hydrochloride (AEBSF, 0.5mM) for 30 min in culture medium before lysis or alternatively 2×10^6^ cells were prepared directly in 100μL NuPAGE LDS sample buffer (1x in RIPA buffer plus inhibitors) and heated at 70°C for 10 min. Where indicated, the proteasome inhibitor MG132 was added to the culture medium at a concentration of 0.25µM for 2h. Chromatin-bound proteins were isolated from subcellular fractions according to BioProtocol 8-9-3035 (Gillotin, 2018). In brief, 6-10×10^6^ myeloid cells were pre-treated with AEBSF as above, then washed with ice-cold PBS and centrifuged at 4°C, followed by sequential extraction of cytoplasmic proteins in E1 buffer (50mM HEPES-KOH, 140mM NaCl, 1mM EDTA, 10% glycerol, 0.5% NP-40, 0.25% Triton X-100, 0.5mM TCEP), nuclear proteins in E2 buffer (10mM Tris-HCl, 200mM NaCl, 1mM EDTA, 05mM EGTA) and chromatin bound proteins in E3 buffer (500mM Tris-HCl, 500mM NaCl). All lysates were sonicated in a Bioruptor sonicating water bath at 4°C, with 5min, 30sec on/off pulses on power high. Protein concentrations were measured with Pierce BCA assay kit and, unless otherwise indicated, 25μg protein extract was separated on pre-cast NuPAGE™ BOLT 8% Bis-Tris polyacrylamide gels (Invitrogen) and transferred onto Immobilon-P PVDF membranes (Merck Millipore). Membranes were blocked with 5% milk in TBST and incubated overnight with rat monoclonal anti-mouse RECQL4 antibody (clone 3B10, made by WEHI Antibody Services, Melbourne) (Castillo-Tandazo *et al*., 2019), mouse anti-β-Actin (Sigma Aldrich, A1978), rabbit anti-Histone H3 (Abcam, ab1719) or rabbit anti-SC-35 (Abcam, ab204916). Membranes were then probed with HRP-conjugated goat anti-rat (Thermo Fisher Scientific, 31470) or anti-mouse (Thermo Fisher Scientific, 31444) secondary antibodies and visualized using ECL Prime Substrate (Amersham). Images were acquired on X-ray film (Fujifilm) or digitally with the iBright Imaging System (Thermo Fisher Scientific) and analysed using ImageJ or iBright software.

### Cell cycle flow cytometry

For DNA replication analysis, cells were treated with 10μM EdU (Life Technologies) for 30min at 37°C. Cells were then harvested, washed in PBS with 1% BSA, and fixed with 2% paraformaldehyde for 15 min at room temperature in the dark. Cells were washed once with PBS 1% BSA and stored short-term at 4°C. To detect incorporated EdU, cells were processed as described by the manufacturer (EdU labelling kit, Life Technologies) using Alexa Fluor 488 Azide or Alexa Fluor 647 Azide. Cells were subsequently stained with anti-Ki-67 PECy7 antibody (eBioscience 25-5698-82) for 30 min, then washed with PBS + 1% BSA and resuspended in 1µg/mL DAPI (Molecular Probes). The various stages of the cell cycle were analysed by flow cytometry as described (Vignon *et al*, 2013).

MCM loading was determined according to Matson *et al*. (Matson & Cook, 2020; Matson *et al*, 2017). In short, cells were pulsed with EdU as above, pre-extracted with CSK buffer (10mM PIPES pH7.0, 300mM sucrose, 100mM NaCl, 3mM MgCl_2_) + 0.5% (V/V) Triton X-100 + protease/phosphatase inhibitors for 5 min on ice, followed by fixation in 4% PFA and washing with PBS 1% BSA. EdU was detected as above and cells were then incubated with mouse anti-MCM3 antibody (Santa Cruz sc-39080) in 0.1% NP40 (Sigma-Aldrich) - PBS 1% BSA. After secondary antibody-staining with anti-mouse Alexa Fluor 594 F(ab’)2 fragments (Cell Signaling Technology #8890), cells were kept in 0.1% NP40 - PBS 1% BSA supplemented with DAPI (1ug/mL) and RNase A (100ug/mL). Cells were analysed by flow cytometry using the FACS BD LSR II Fortessa system and FlowJo software version 10 (BD Bioscience).

### Peripheral blood analysis

Peripheral blood was analysed on a hematological analyser (Sysmex KX-21N, Roche Diagnostics). For flow cytometric analysis, red blood cells were lysed using a red blood cell lysis buffer (150mM NH_4_Cl, 10mM KHCO_3_, 0.1mM Na_2_EDTA, pH7.3).

### Flow cytometry analysis of mouse tissues

Femurs were flushed (2 femurs in 2mL PBS/2%FBS), spleens (5mL PBS/2%FCS) and thymus (2mL PBS/2%FCS) crushed, and single cell suspensions were prepared in PBS containing 2% FBS. Antibodies against murine B220 (APC-eFluor780), CD11b/Mac-1 (PE), Gr1 (PE-Cy7), F4/80 (APC), CD4 (eFluor450) and CD8a (PerCP-Cy5.5), Ter119 (PE), CD71 (APC), CD44 (PE-Cy7), Sca-1 (PerCP-Cy5.5), c-Kit (APC-eFluor780), CD150 (PE), CD48 (PE-Cy7), CD34(eFluor660), CD16/32 (eFluor450) and biotinylated antibodies (CD2, CD3e, CD4, CD5, CD8a, B220, Gr-1, CD11b/Mac1, Ter-119) were used. The biotinylated antibodies were detected with streptavidin-conjugated Brilliant Violet-605 (Liddicoat *et al*, 2015; Singbrant *et al*, 2011; Smeets *et al*., 2014). 30,000-500,000 cells were acquired on a BD LSRII Fortessa and analysed with FlowJo software Version 9 or 10.0 (Treestar).

### Global Protein Stability (GPS) assays

The empty GPS reporter construct and constructs containing USP49, PPP1R15A, NS3, NS3GA and NS8 were a kind gift from Prof. Hsueh-Chi S. Yen (IMB, Academia Sinica, Taipei, Taiwan) and have been described before (Lin *et al*, 2018; Yen & Elledge, 2008; Yen *et al*, 2008). The original pDEST-GPS Gateway vector (Huang *et al*, 2023) was adapted to allow for standard cloning by replacing a 1721bp BsrG1 fragment with a BsrG1-NotI-co*Cenpw*-XhoI-BsrGI gene fragment. The cDNA encoding the predicted last 172aa of the recombined/deleted *Recql4* allele was synthesised from *Recql4^Δ/Δ^ sgKlhdc3* derived mRNA and cloned between NotI and XhoI in-frame with the GFP. Three clones containing the correct C-terminal sequence of this short product (S4, S5 and S6) were transduced individually in HEK293T WT and KLHDC3 KO cells. Human codon-optimised *KLHDC3* was obtained from Twist Bioscience as a gene fragment with an N-terminal 3xFlag tag and cloned into the pAIP (pLenti SFFV IRES puromycin, Plasmid #74171, Addgene) and pLenti SFFV SV40 blasticidin vectors. HEK293T KLHDC3 WT or KO cells were transduced with GPS constructs and selected with 1µg/mL puromycin. Where indicated, KLHDC3 was re-or over-expressed by transduction with pLenti SFFV 3xFlag-coKLHDC3 and selection with 10µg/mL blasticidin. Protein stability was visualised by microscopy on an inverted fluorescence microscope (Olympus IX81) using cellSens (Olympus Lifescience) software for image acquisition and analysed by flow cytometry using a BD LSRIIFortessa for acquisition and FlowJo software Version 10.0 (BD Biosciences) to determine the ratio of GFP/DsRed for each reporter construct (Lin *et al*., 2018).

### DNA replication timing profiling by whole genome sequencing

Extraction of genomic DNA was performed using the Gentra Puregene kit (Qiagen) following the manufacturer’s instructions. The purified DNA was then sent to BGI Tech Solutions (Hong Kong, China) for library preparation and Whole Genome Sequencing (150PE, DNBSEQ). DNA replication timing profiles across the genome were obtained using the Timing Inferred from Genome Replication (TIGER) methodology as described (Bracci *et al*, 2023; Koren *et al*, 2021). We assessed the data in 10kb windows and with the remaining settings as default parameters.

### Datasets and sequencing files

Screen results datasets are contained in the supplemental files and related sequencing files have been deposited in GEO under accession numbers GSE273174 (screen in Figure 1), GSE273006 (synthetic lethal screen) and GSE272599 (whole genome sequencing dataset used to calculate replication timing).

## Statistical analysis

To determine statistical significance, log-rank tests, t-tests and ordinary one-way ANOVA with multiple comparison correction were conducted in GraphPad Prism software version 9 or 10 (GraphPad; San Diego, CA, USA). Statistical significance is indicated as: *p<0.05; **p<0.01; ***p<0.001 and****p<0.0001, and data are presented as mean±SEM unless otherwise indicated. The number of samples used for each experiment is indicated in the figure panels or described in the corresponding figure legends.

## Supplemental Information Datasets

### Dataset S1. Initial genome-wide CRISPR knock-out screen (described in Figure 1)

Related datalinks:

The following secure token has been created to allow review of record GSE273174 while it remains in private status:

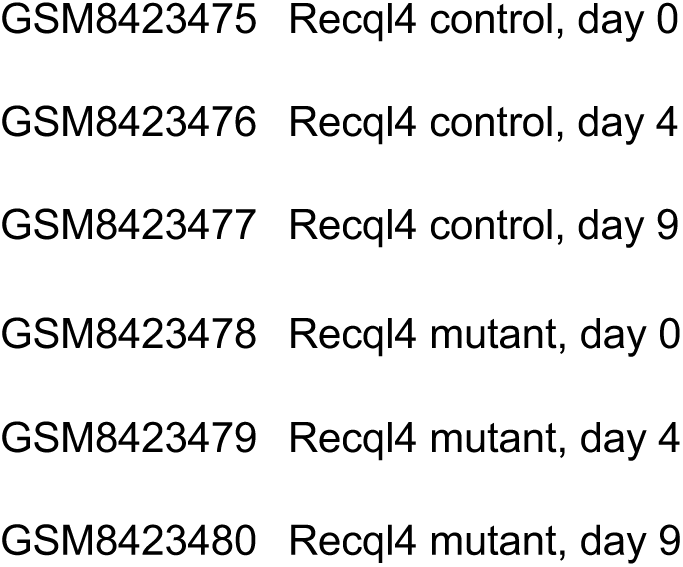

### Dataset S2. Synthetic lethal genome-wide CRISPR knock-out screen (described in Figure 4)

Related datalinks:

The following secure token has been created to allow review of record GSE273006 while it remains in private status:

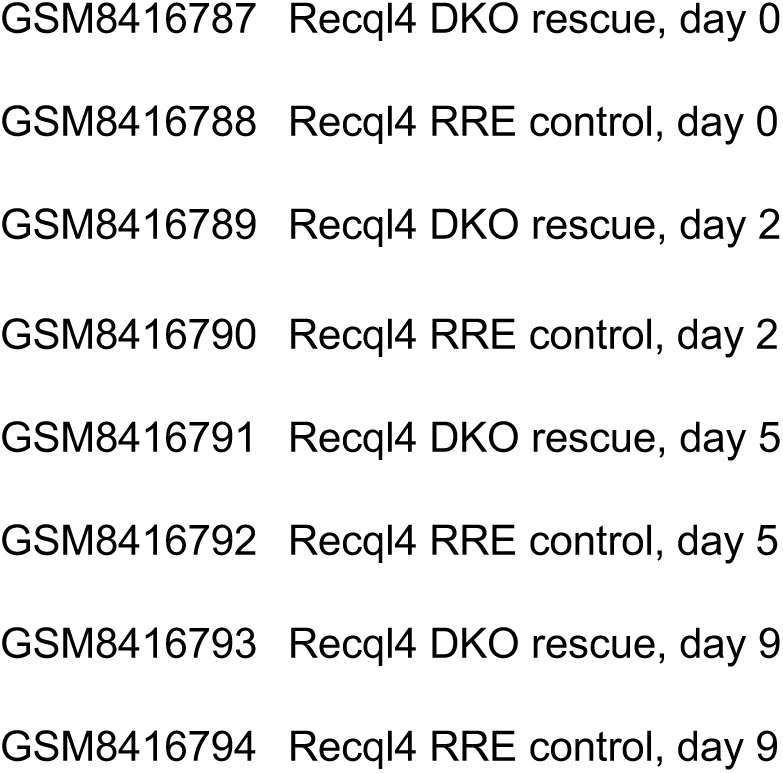

### Whole Genome sequencing dataset used to calculate replication timing

The following secure token has been created to allow review of record GSE272599 while it remains in private status:

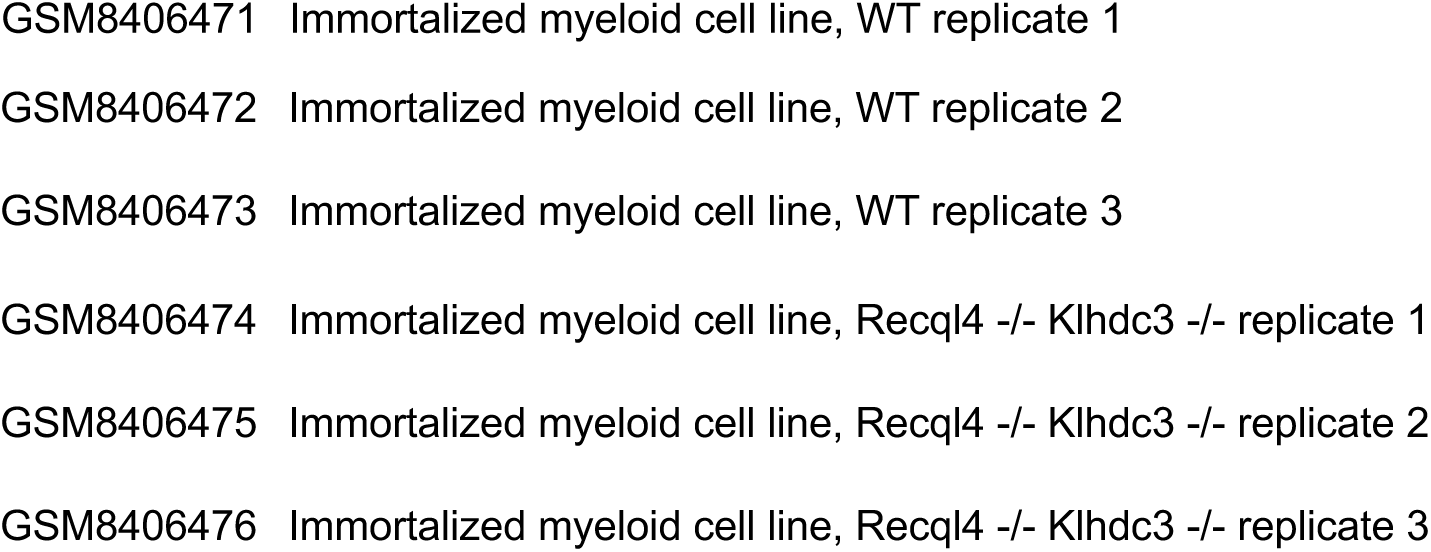

### Reagents

#### sgRNA sequences used in this study

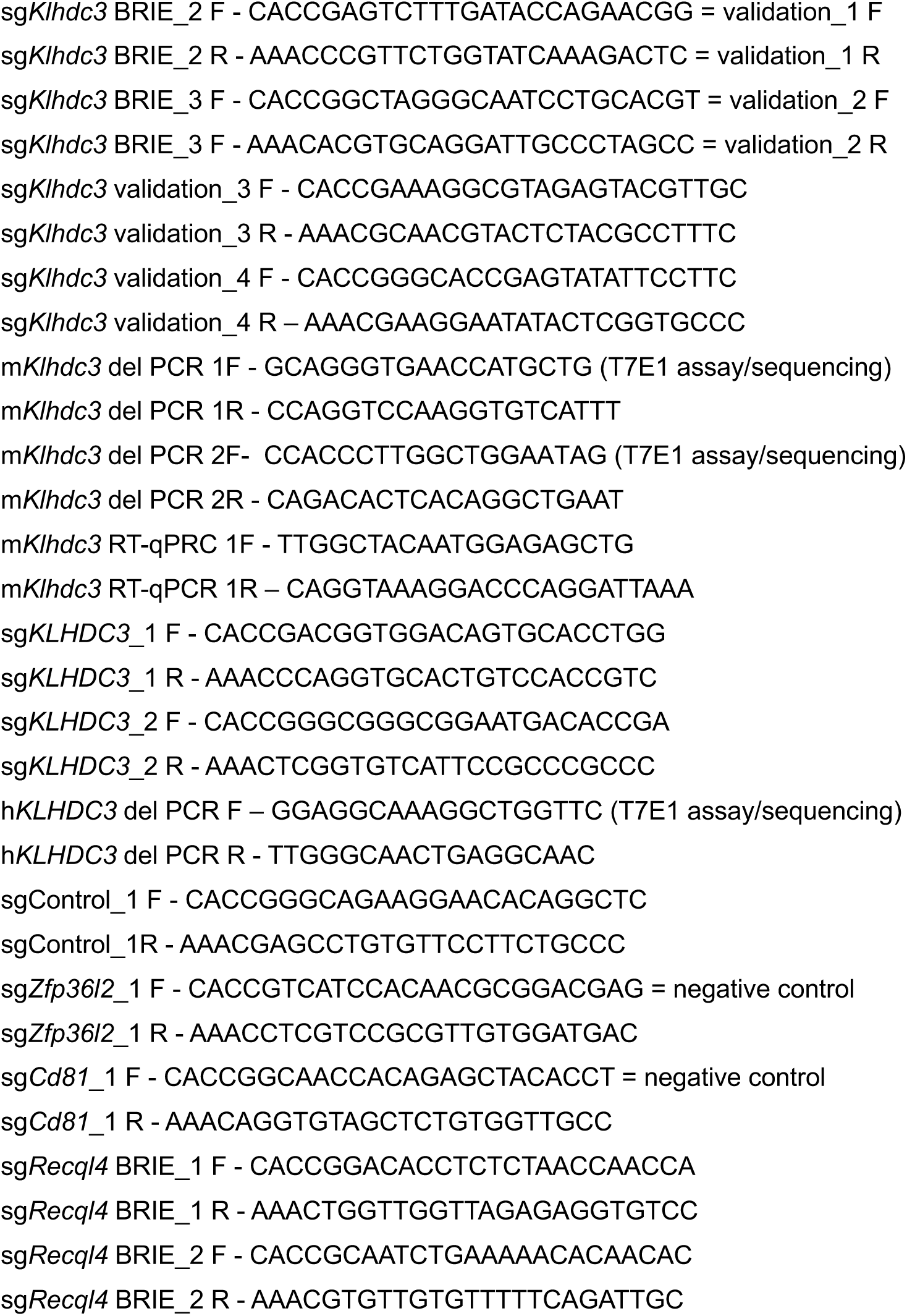
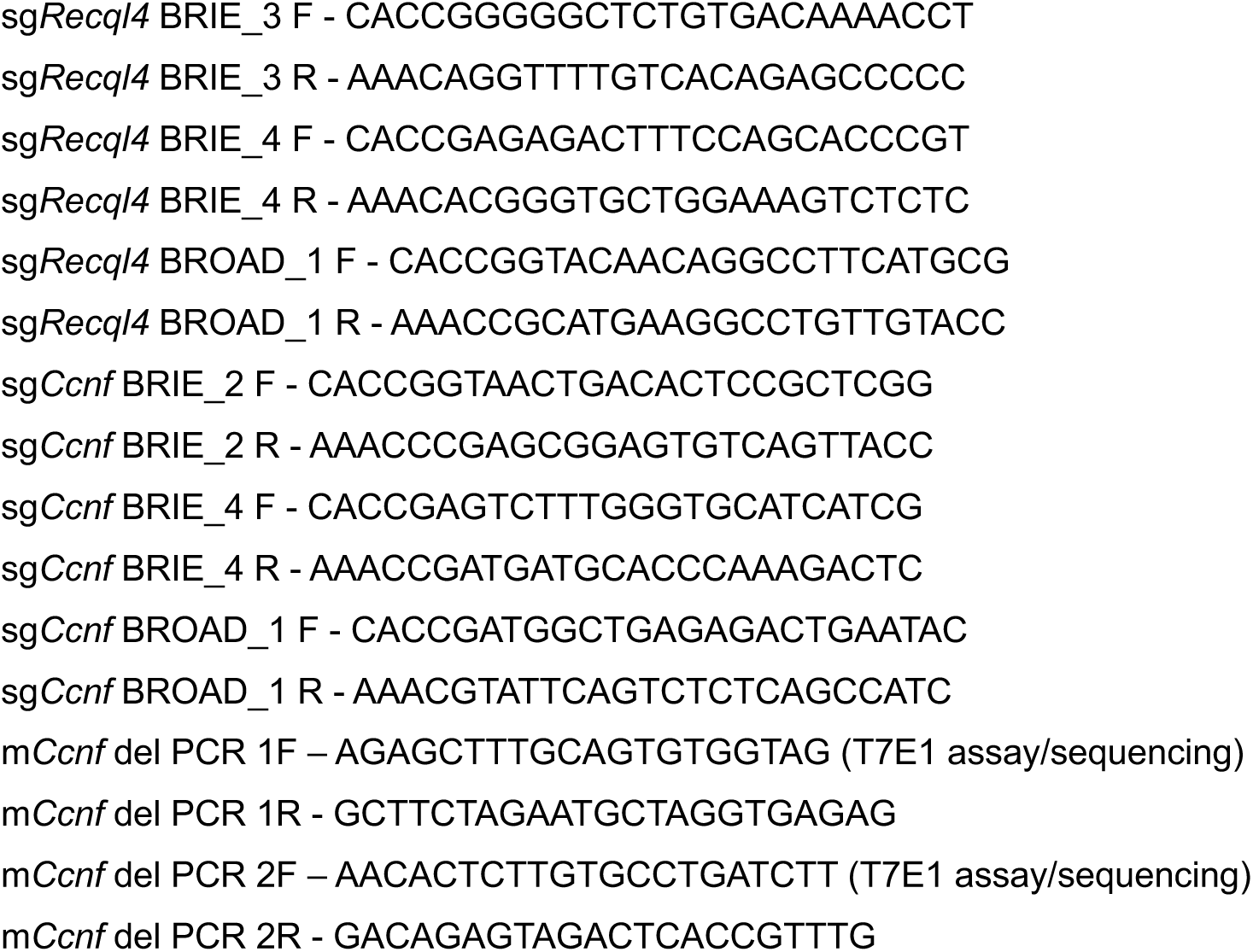

### Genotyping primers (all 5’-3’) Recql4

WT ACAGCAACAGAACAGCAACTACG;

floxed CACTCTAGAAGAGGGAGTCAGATGG;

deleted CGCGCGAAAGCTGAGGAGTT;

product size: WT = 165bp, floxed =325bp; deleted =256bp; reference Smeets et al.(Smeets *et al*., 2014)

### Klhdc3

WT GCATGGTGGAACAAGAGACTTT;

floxed GCCAAGAGCAGAGAGATGGG;

deleted CTGTGGCCCAGGGGATGTTA;

product size: WT = 306bp, floxed =386bp; recombined = 552bp; germ-line deleted =451bp

**R26-CreER** (RRID:IMSR_JAX:008463):

Primer 1 AAAGTCGCTCTGAGTTGTTAT;

Primer 2 CCTGATCCTGGCAATTTCG; Primer 3 GGAGCGGGAGAAATGGATATG;

product size: WT = 650bp, KI =825bp

### Recql4 G522Efs and R347* alleles

The presence of the G522Efs (Castillo-Tandazo *et al*., 2019) and R347* (Castillo-Tandazo *et al*., 2019) mutations was determined by KASP (competitive allele specific PCR) technology (LGC) with custom designed (G522Efs) or facility provided (R347X primer: 5’-GAAGGTGACCAAGTTCATGCTAAAGCGTTTGTTTTTCATGTTGAGTCG-3’, 5’-GAAGGTCGGAGTCAACGGATTCAAAGCGTTTGTTTTTCATGTTGAGTCA-3’, reverse primer 5’-GCTTCCCTAGACAGAGGGAACTATA-3’) sequences according to manufacturer instructions.

## Supplemental Figure Legends

**Figure S1.**
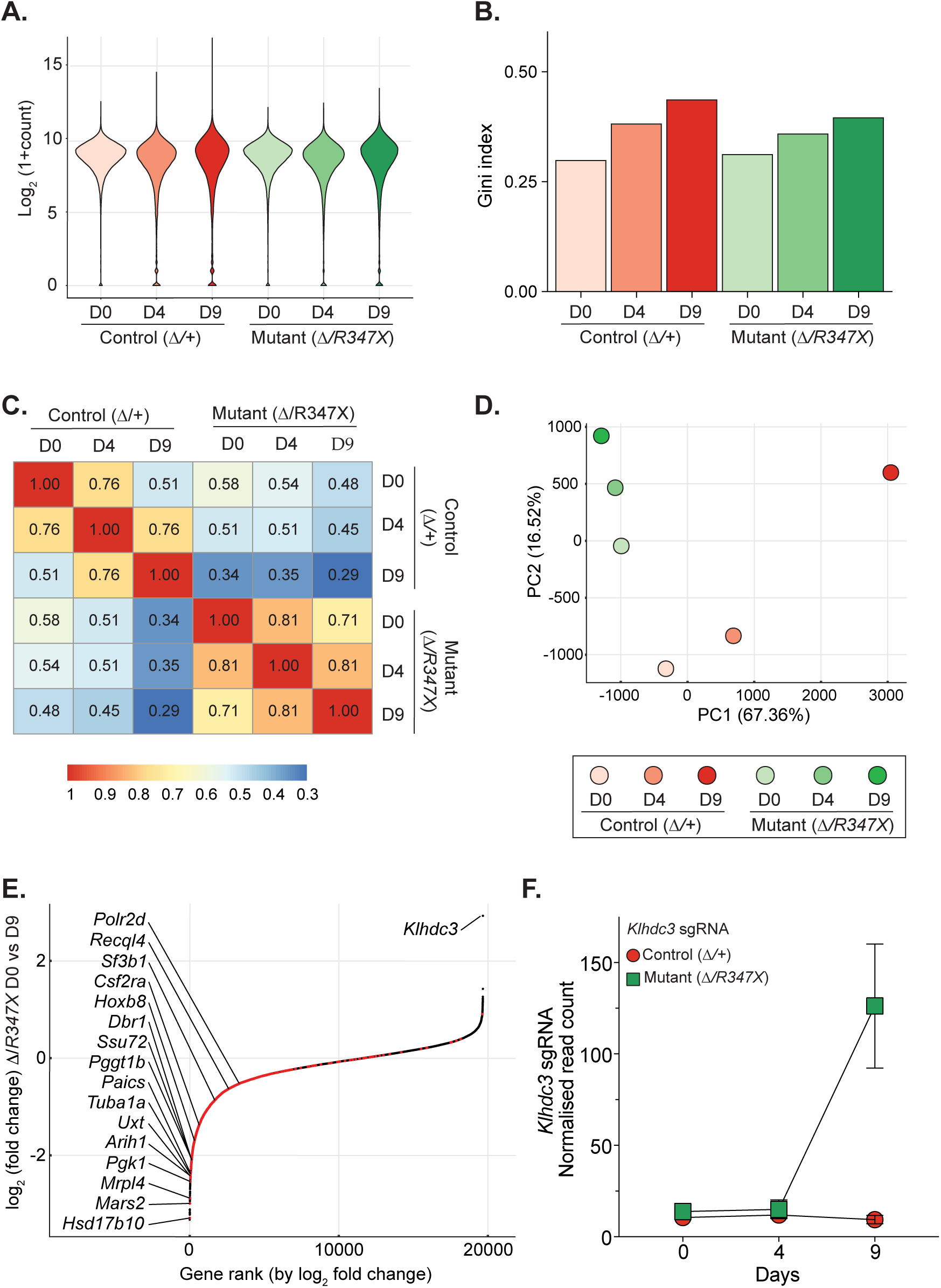
Loss of function rescue screen metrics. A. sgRNA count distribution. Violin Plot showing sgRNA frequencies at day 0, 4, and 9 and control (fl/+) and mutant (fl/R347X) cell lines B. Gini index computed from normalised sgRNA counts for all cell line replicates. C. Heatmap showing sgRNA-level correlation between timepoints of control and mutant cell lines. D. Principle component analysis (PCA) of sgRNA-sequencing results from day 0, 4, and 9 of control and mutant cell lines E. Essential gene depletion. sgRNAs of R347 D9 vs D0 were ranked by log2 fold change, showing loss of representation of known and predicted essential genes. F. Normalised sgRNA counts for the four *Klhdc3* guides in the library.

**Figure S2.**
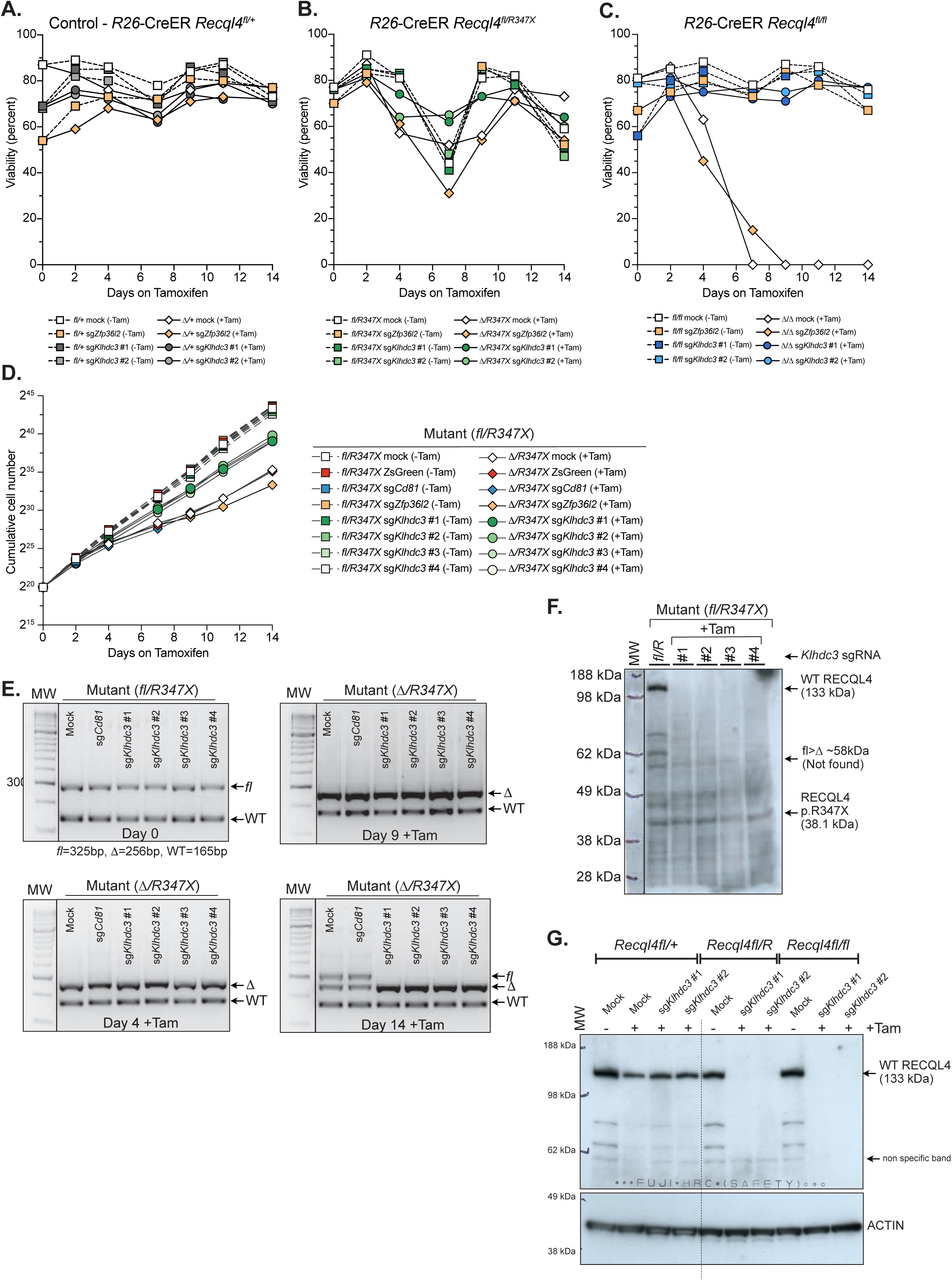
Additional validation that loss of *Klhdc3* rescues the proliferative defect of both *Recql4* point mutant and deficient myeloid cells. A. Effect of loss of *Klhdc3* on control cells (*R26*-CreER *Recql4^fl/+^*; become Δ*/+* cell following tamoxifen treatment applied at Day 0) viability (related to Figure 1C). B. Effect of loss of *Klhdc3* on *Recql4* p.R347X only expressing cells (*R26*-CreER *Recql4^fl/R347X^*; become Δ*/R347X* cells following tamoxifen treatment applied at Day 0) viability (related to Figure 1D). C. Effect of loss of *Klhdc3* on *Recql4* deficient cells (*R26*-CreER *Recql4^fl/fl^*; become Δ*/*Δ cells following tamoxifen treatment applied at Day 0) viability (related to Figure 1E). D. Proliferation assay showing that four sgRNAs against *Klhdc3* (two from the BRIE library and two new guides) rescue *Recql4^R347X^*point mutant cells (circles). Results were compared to non-tamoxifen treated cells (squares), and sgRNA targeting *Cd81* (a non-essential cell surface marker), sg*Zfp36l2* (a guide depleted in the mutant but not in the control), ZsGreen (a non-targeting virus control), and a mock (non-infected) control (diamonds). E. Genomic PCR showing successful recombination and stable deletion of the floxed *Recql4* allele after addition of tamoxifen in *Klhdc3 Recql4* R347X double mutant cells. F. Western blot of full-length WT and truncated R347X mutant RECQL4 protein in the non-tamoxifen treated control and tamoxifen-treated *Klhdc3* sgRNA mutant cells (Day 9 Post-Tam). G. Western blot of control and tamoxifen treated *Recql4^fl/+^* (heterozygous control after tamoxifen treatment), *Recql4^R347X/fl^* and *Recql4^fl/fl^* myeloid cells probing for RECQL4 or ACTIN as indicated. Treatment as indicated (Mock = non-infected or sg*Klhdc3* #1 or #2).

**Figure S3.**
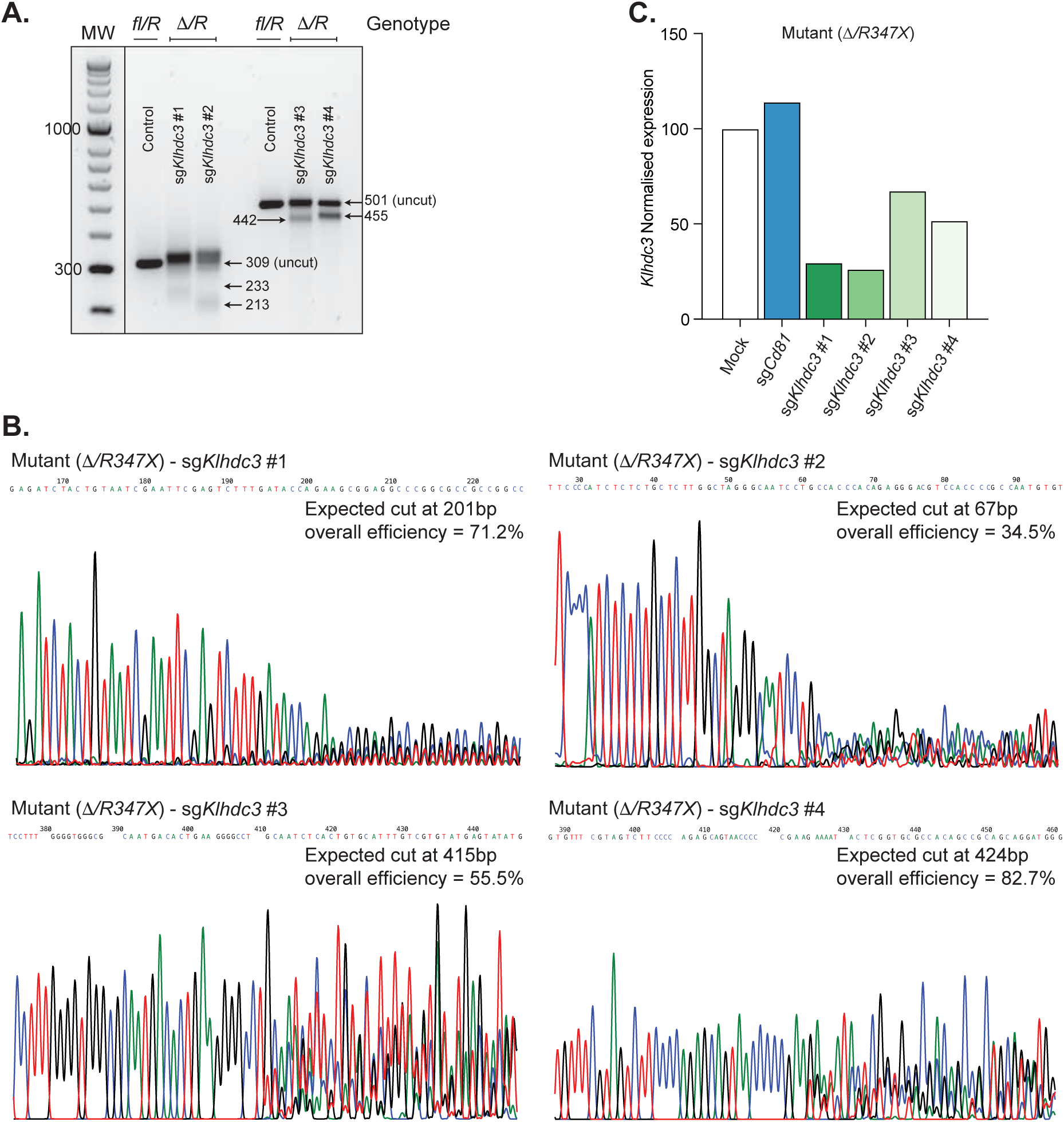
Validation of *Klhdc3* targeting by sgRNA. A. T7 endonuclease digestion of heteroduplex PCR products amplified from the genomic DNA surrounding the sgRNA target sites, showing cleavage of mismatched DNA at sites of indels ≥2 bases in a mixed population of *Klhdc3* gRNA targeted mutant (R347X) cells (Day 9 Post-Tam). The sizes of the predicted cleavage products are labelled in the figure. B. TIDE Quantitative Sanger sequence trace analysis of amplified genomic DNA regions surrounding the expected break site of individual *Klhdc3* sgRNAs in a mixed population of mutant (R347X) cells (Day 9 Post-Tam). All cells were subsequently cloned and sequenced to ensure homozygous mutations were present. C. Quantitative real-time RT-PCR analysis of *Klhdc3* RNA expression in mock and tamoxifen-treated *sgKlhdc3 Recql4 R347X* double mutant cells (Day 9 Post-Tam). (Expression relative to *Gapdh* and normalised to mock).

**Figure S4.**
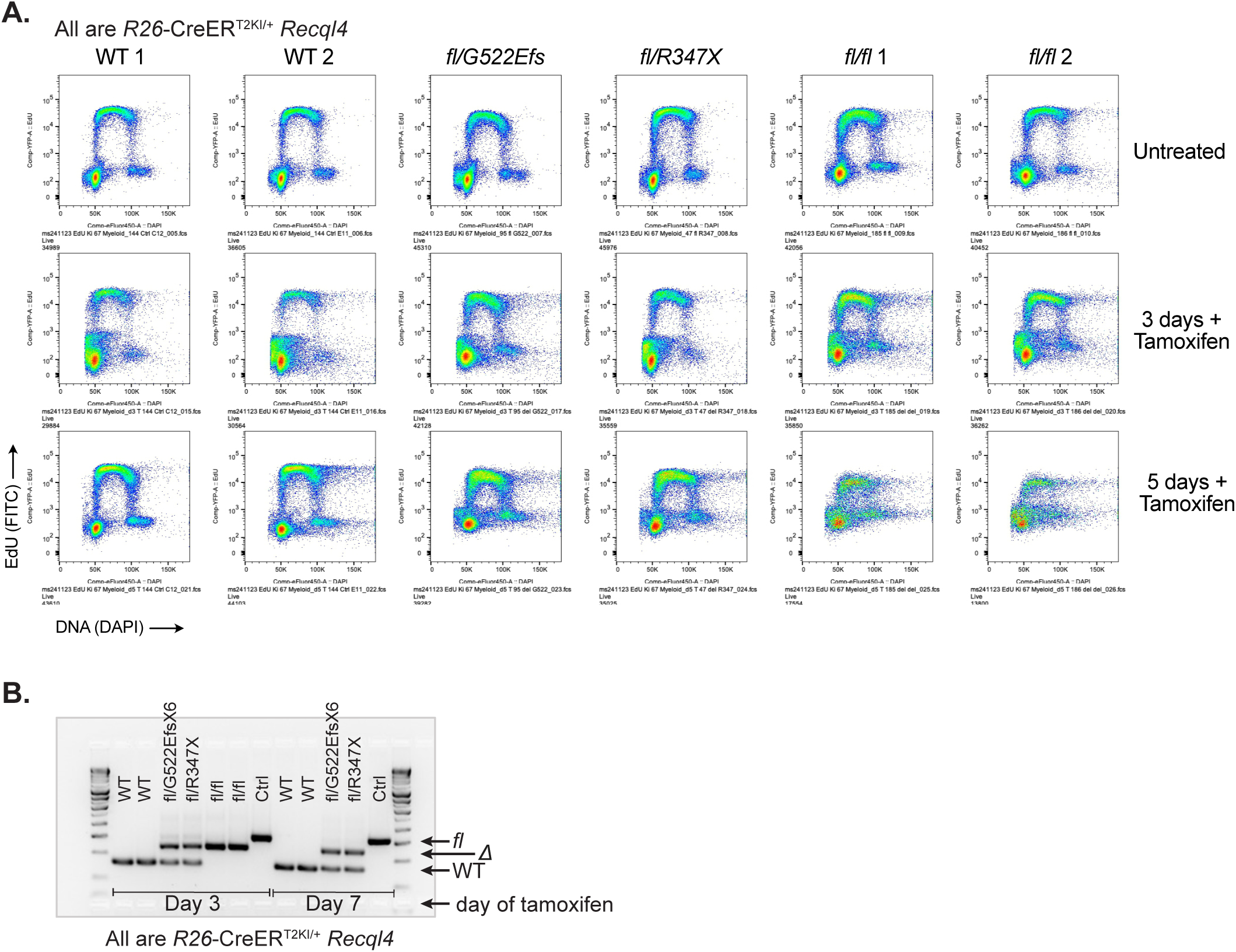
Cell cycle kinetics of WT, point mutant and *Recql4* deficient myeloid cells. A. Representative flow cytometry plots assessing DNA replication rates following pulse labelling of asynchronous cultures of myeloid cells with EdU incorporation and DAPI (DNA stain). Genotypes assessed as *Recql4* wild-type (WT), *Recql4^fl/G522Efs^*, *Recql4^fl/R347X^* and *Recql4^fl/fl^*. All cell lines are *R26*-CreER^T2ki/+^ and *Klhdc3* wild-type. Day after tamoxifen addition as indicated. B. Genomic PCR showing recombination of the floxed *Recql4* allele at the indicated days after the addition of tamoxifen in each respective genotype. All cells are *Klhdc3^+/+^*.

**Figure S5.**
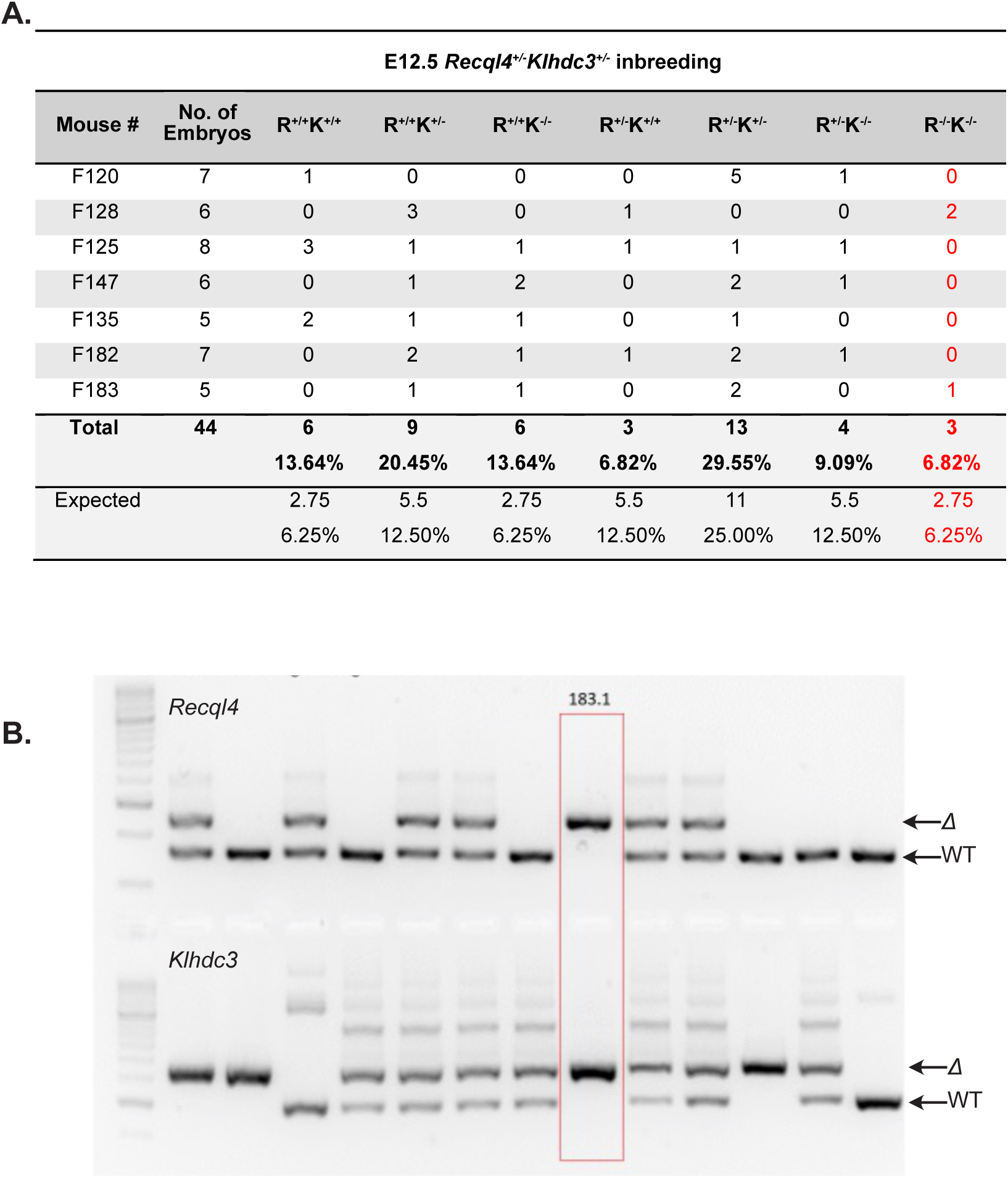
Loss of *Klhdc3* extends the survival of *Recql4* deficient embryos *in vivo*. A. Recovery of indicated genotypes at embryonic day 12.5 (E12.5) from inbreeding of *Recql4^+/-^ Klhdc3^+/-^* breeding pairs. Previous analysis demonstrated that the *Recql4^-/-^* embryos were lethal prior to E10.5 (specific time point prior to this not determined). B. Genomic PCR showing demonstrating recovery of a homozygous embryo.

**Figure S6.**
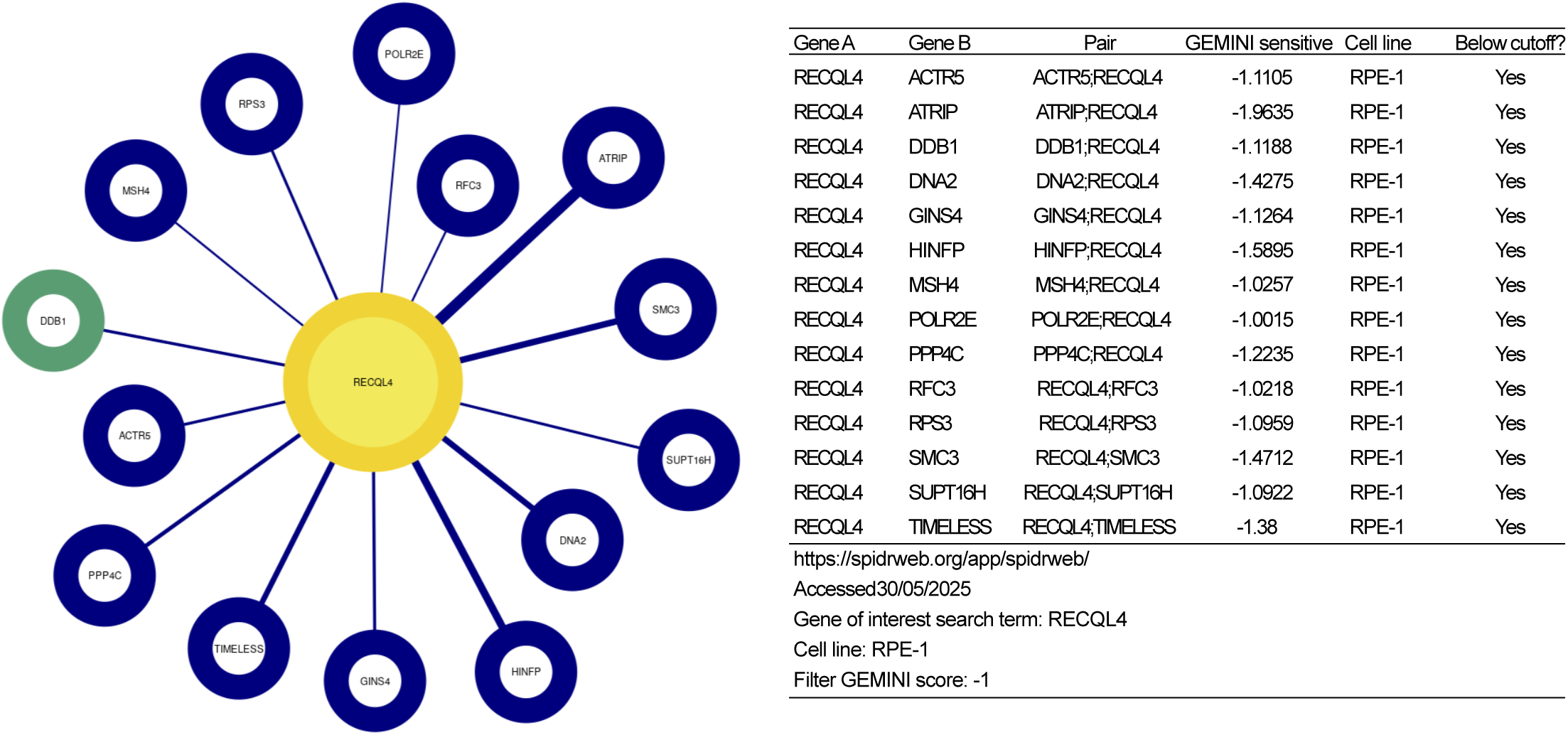
Synthetic lethal interactions with RECQL4 in human RPE-1 cells. Data from spidrweb.org/app/spidrweb/; SPIDRweb: Comprehensive Interrogation of Synthetic Lethality in the DNA Damage Response Published in *Nature*: https://doi.org/10.1038/s41586-025-08815-4.

**Figure S7.**
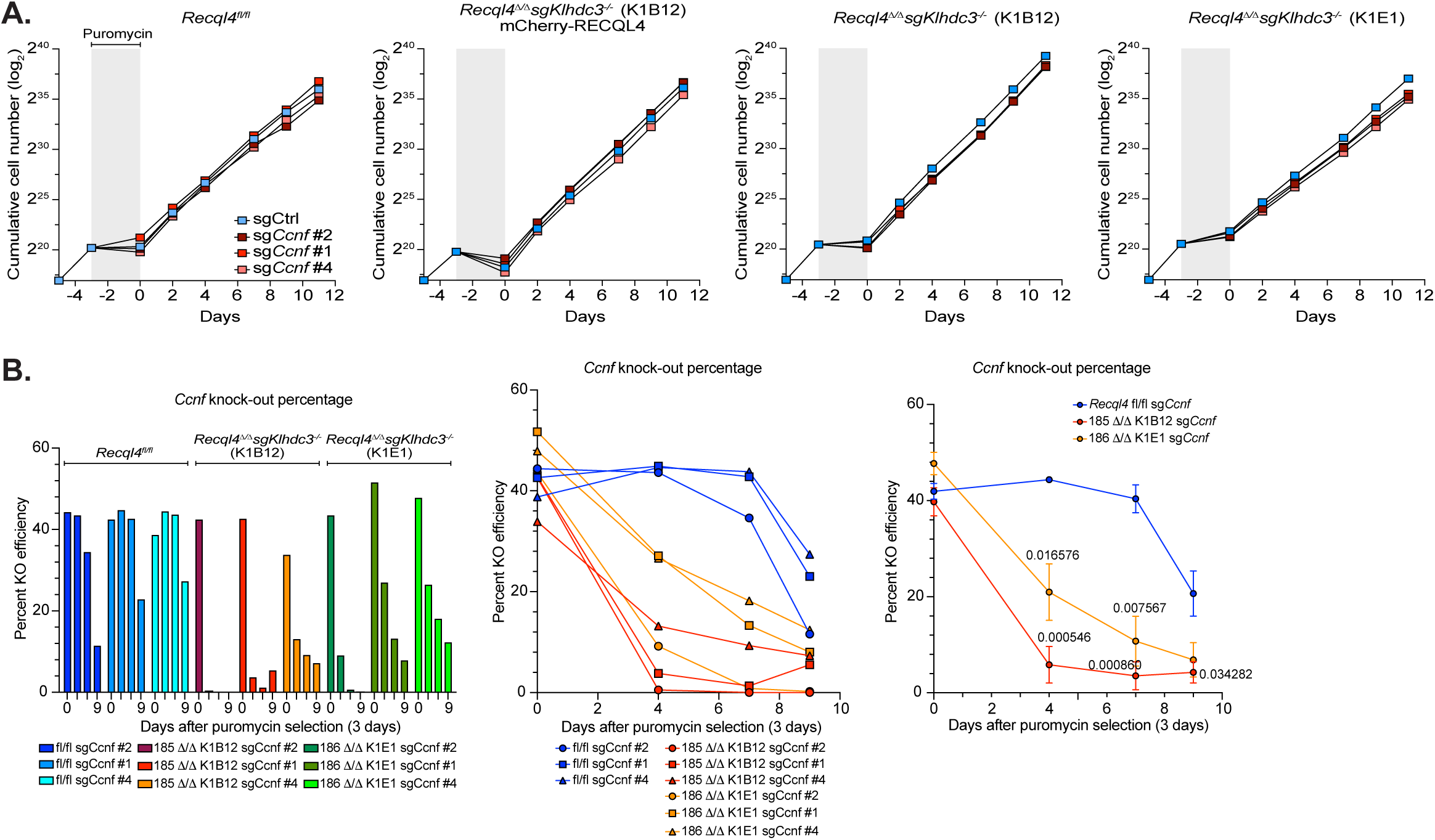
Validation that loss of Cyclin F is synthetic lethal with Recql4 deficiency. A. Proliferation curves of sgCtrl and sg*Ccnf*-targeted cell lines of the indicated genotypes. Grey shaded area indicates puromycin selection. *Recql4^1/1^* sg*Klhdc3* K1B12 and *Recql4^1/1^* sg*Klhdc3* K1E1 are independently targeted and isolated clones. B. The knockout efficiency of *Ccnf* was measured through Sanger Sequencing and analysed by TIDE at each time point in each genotype as indicated. Data shown as each sample individually and as the mean knockout efficiency of three sg*Ccnf* guides +/-SEM (right panel). Statistical analysis was done using multiple unpaired t-tests with significant p-values listed.

**Figure S8.**
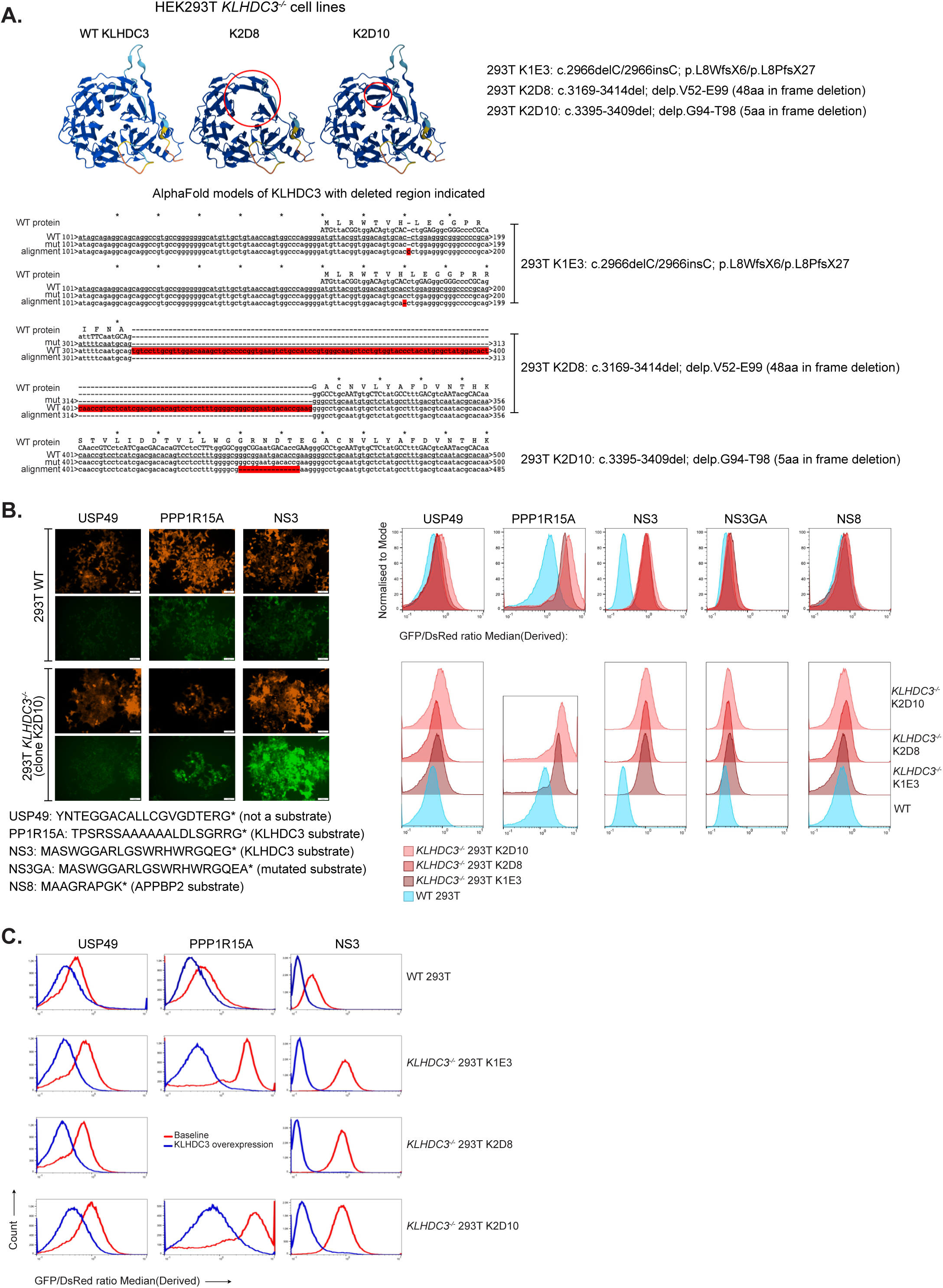
Generation of KLHDC3^-/-^ 293T cells. A. Schematic and analysis of 3 independent *KLHDC3^-/-^* 293T cell lines. Each line was confirmed as a homozygous mutant. B. Validation of KLHDC3 deficiency using GPS reporter assay by either live cell fluorescent microscopy of flow cytometry. Representative images of GPS reporters for USP49, PPP1R15A and NS3 (all KLHDC3 substrates) and NS3GA and NS8 (not KLHDC3 substrates) by live cell fluorescent imaging in WT and a *KLHDC3^-/-^*293T cell (clone K2D10). Scale bar represents 50*μ*m. Representative flow cytometric analysis of the GFP and DsRed expression in 3 independently generated *KLHDC3^-/-^* 293T cells compared to KLHDC3 WT 293T cells. Stabilisation of GFP is indicated by the shift in GFP/DsRed derived median. C. Re-expression of KLHDC3 in the *KLHDC3^-/-^*293T cells leads to loss of GFP signal for the known KLHDC3 substrates USP49, PP1R15A and NS3. The baseline GFP/DsRed derived median is in red, the KLHDC3 over-expressing samples are in blue. Note the left shift in the WT KLHDC3 re-expressing *KLHDC3^-/-^* 293T cells indicative of destruction of the GFP-fusion protein.

**Figure S9.**
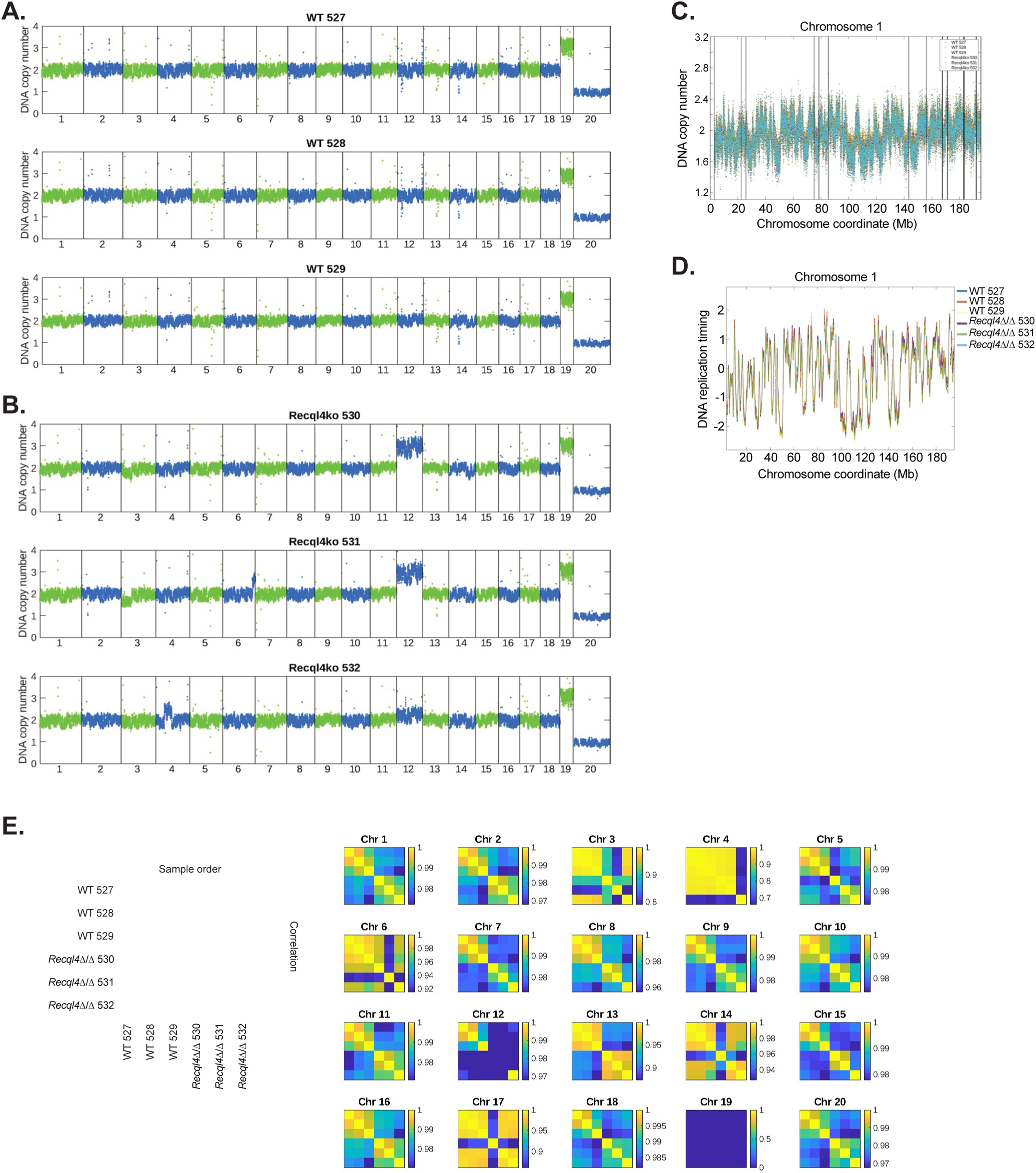
Individual whole genome DNA copy number analysis for the myeloid cell lines used for DNA replication timing inference. A. DNA copy number across all chromosomes for WT myeloid cell lines. B. DNA copy number across all chromosomes for *Recql4^1/1^* sg*Klhdc3* myeloid cell lines. C. Whole genome sequencing was used to infer DNA replication timing across the genome in 3 control and 3 *Recql4^1/1^* sg*Klhdc3* cell lines (WT: 144 *Recql4* +/+ LCr. Hygro Ctrl C12, E12 and H11 and *Recql4* Δ/Δ: 185 *Recql4* Δ/Δ LCr. Hygro *sgKlhdc3* K2C12 and K1B12 and 186 *Recql4* Δ/Δ LCr. Hygro sg*Klhdc3* K1E1). Representative example from each sample is plotted individually for Chromosome 1. D. Representative example of smoothed plots of DNA replication timing across chromosome 1 for each cell line. E. Correlation analysis of DNA replication timing of each chromosome across the genome with the scale for each individual chromosome. Note that missing data, due to filtering of specific chromosomes in some samples, appears as dark blue in the correlation matrices.

## References

Bhowmick R, Hickson ID, Liu Y (2023) Completing genome replication outside of S phase. Mol Cell 83: 3596–3607

Bhowmick R, Minocherhomji S, Hickson ID (2016) RAD52 Facilitates Mitotic DNA Synthesis Following Replication Stress. Mol Cell 64: 1117–1126

Bracci AN, Dallmann A, Ding Q, Hubisz MJ, Caballero M, Koren A (2023) The evolution of the human DNA replication timing program. Proc Natl Acad Sci U S A 120: e2213896120

Brinkman EK, Kousholt AN, Harmsen T, Leemans C, Chen T, Jonkers J, van Steensel B (2018) Easy quantification of template-directed CRISPR/Cas9 editing. Nucleic Acids Res 46: e58

Castillo-Tandazo W, Frazier AE, Sims NA, Smeets MF, Walkley CR (2021) Rothmund-Thomson Syndrome-Like RECQL4 Truncating Mutations Cause a Haploinsufficient Low-Bone-Mass Phenotype in Mice. Mol Cell Biol 41: e0059020

Castillo-Tandazo W, Smeets MF, Murphy V, Liu R, Hodson C, Heierhorst J, Deans AJ, Walkley CR (2019) ATP-dependent helicase activity is dispensable for the physiological functions of Recql4. PLoS Genet 15: e1008266

Chu WK, Hickson ID (2009) RecQ helicases: multifunctional genome caretakers. Nat Rev Cancer 9: 644–654

Clijsters L, Hoencamp C, Calis JJA, Marzio A, Handgraaf SM, Cuitino MC, Rosenberg BR, Leone G, Pagano M (2019) Cyclin F Controls Cell-Cycle Transcriptional Outputs by Directing the Degradation of the Three Activator E2Fs. Mol Cell 74: 1264–1277 e1267

Conant D, Hsiau T, Rossi N, Oki J, Maures T, Waite K, Yang J, Joshi S, Kelso R, Holden K et al. (2022) Inference of CRISPR Edits from Sanger Trace Data. CRISPR J 5: 123–130

Cvetkovic MA, Passaretti P, Butryn A, Reynolds-Winczura A, Kingsley G, Skagia A, Fernandez-Cuesta C, Poovathumkadavil D, George R, Chauhan AS et al. (2023) The structural mechanism of dimeric DONSON in replicative helicase activation. Mol Cell 83: 4017–4031 e4019

Doench JG, Fusi N, Sullender M, Hegde M, Vaimberg EW, Donovan KF, Smith I, Tothova Z, Wilen C, Orchard R et al (2016) Optimized sgRNA design to maximize activity and minimize off-target effects of CRISPR-Cas9. Nat Biotechnol 34: 184–191

Durinck S, Bullard J, Spellman PT, Dudoit S (2009a) GenomeGraphs: integrated genomic data visualization with R. BMC Bioinformatics 10: 2

Durinck S, Spellman PT, Birney E, Huber W (2009b) Mapping identifiers for the integration of genomic datasets with the R/Bioconductor package biomaRt. Nat Protoc 4: 1184–1191

Evrin C, Alvarez V, Ainsworth J, Fujisawa R, Alabert C, Labib KP (2023) DONSON is required for CMG helicase assembly in the mammalian cell cycle. EMBO Rep 24: e57677

Fielden J, Siegner SM, Gallagher DN, Schroder MS, Dello Stritto MR, Lam S, Kobel L, Schlapansky MF, Jackson SP, Cejka P et al (2025) Comprehensive interrogation of synthetic lethality in the DNA damage response. Nature 640: 1093–1102

Gillotin S (2018) Isolation of Chromatin-bound Proteins from Subcellular Fractions for Biochemical Analysis. Bio Protoc 8: e3035

Hart T, Tong AHY, Chan K, Van Leeuwen J, Seetharaman A, Aregger M, Chandrashekhar M, Hustedt N, Seth S, Noonan A et al (2017) Evaluation and Design of Genome-Wide CRISPR/SpCas9 Knockout Screens. G3 (Bethesda) 7: 2719–2727

Hashimoto Y, Sadano K, Miyata N, Ito H, Tanaka H (2023) Novel role of DONSON in CMG helicase assembly during vertebrate DNA replication initiation. EMBO J 42: e114131

Heraud-Farlow JE, Taylor SR, Chalk AM, Escudero A, Hu SB, Goradia A, Sun T, Li Q, Nikolic I, Li JB et al (2024) GGNBP2 regulates MDA5 sensing triggered by self double-stranded RNA following loss of ADAR1 editing. Sci Immunol 9: eadk0412

Hickson ID (2003) RecQ helicases: caretakers of the genome. Nat Rev Cancer 3: 169–178

Huang WC, Yeh CW, Hsu SY, Lee LT, Chu CY, Yen HS (2023) Characterization of degradation signals at protein C-termini. Methods Enzymol 686: 345–367

Ichikawa K, Noda T, Furuichi Y (2002) [Preparation of the gene targeted knockout mice for human premature aging diseases, Werner syndrome, and Rothmund-Thomson syndrome caused by the mutation of DNA helicases]. Nihon Yakurigaku Zasshi 119: 219–226

Jeong HH, Kim SY, Rousseaux MWC, Zoghbi HY, Liu Z (2019) Beta-binomial modeling of CRISPR pooled screen data identifies target genes with greater sensitivity and fewer false negatives. Genome Res 29: 999–1008

Jin W, Liu H, Zhang Y, Otta SK, Plon SE, Wang LL (2008) Sensitivity of RECQL4-deficient fibroblasts from Rothmund-Thomson syndrome patients to genotoxic agents. Hum Genet 123: 643–653

Kingsley G, Skagia A, Passaretti P, Fernandez-Cuesta C, Reynolds-Winczura A, Koscielniak K, Gambus A (2023) DONSON facilitates Cdc45 and GINS chromatin association and is essential for DNA replication initiation. Nucleic Acids Res 51: 9748–9763

Kitao S, Lindor NM, Shiratori M, Furuichi Y, Shimamoto A (1999a) Rothmund-thomson syndrome responsible gene, RECQL4: genomic structure and products. Genomics 61: 268–276

Kitao S, Shimamoto A, Goto M, Miller RW, Smithson WA, Lindor NM, Furuichi Y (1999b) Mutations in RECQL4 cause a subset of cases of Rothmund-Thomson syndrome. Nat Genet 22: 82–84

Koren A, Massey DJ, Bracci AN (2021) TIGER: inferring DNA replication timing from whole-genome sequence data. Bioinformatics 37: 4001–4005

Koren I, Timms RT, Kula T, Xu Q, Li MZ, Elledge SJ (2018) The Eukaryotic Proteome Is Shaped by E3 Ubiquitin Ligases Targeting C-Terminal Degrons. Cell 173: 1622–1635 e1614

Larizza L, Roversi G, Volpi L (2010) Rothmund-Thomson syndrome. Orphanet J Rare Dis 5: 2

Li W, Koster J, Xu H, Chen CH, Xiao T, Liu JS, Brown M, Liu XS (2015) Quality control, modeling, and visualization of CRISPR screens with MAGeCK-VISPR. Genome Biol 16: 281

Liddicoat BJ, Piskol R, Chalk AM, Ramaswami G, Higuchi M, Hartner JC, Li JB, Seeburg PH, Walkley CR (2015) RNA editing by ADAR1 prevents MDA5 sensing of endogenous dsRNA as nonself. Science 349: 1115–1120

Lim Y, Tamayo-Orrego L, Schmid E, Tarnauskaite Z, Kochenova OV, Gruar R, Muramatsu S, Lynch L, Schlie AV, Carroll PL et al (2023) In silico protein interaction screening uncovers DONSON’s role in replication initiation. Science 381: eadi3448

Lin HC, Ho SC, Chen YY, Khoo KH, Hsu PH, Yen HC (2015) SELENOPROTEINS. CRL2 aids elimination of truncated selenoproteins produced by failed UGA/Sec decoding. Science 349: 91–95

Lin HC, Yeh CW, Chen YF, Lee TT, Hsieh PY, Rusnac DV, Lin SY, Elledge SJ, Zheng N, Yen HS (2018) C-Terminal End-Directed Protein Elimination by CRL2 Ubiquitin Ligases. Mol Cell 70: 602–613 e603

Lu H, Davis AJ (2021) Human RecQ Helicases in DNA Double-Strand Break Repair. Front Cell Dev Biol 9: 640755

Lu H, Shamanna RA, de Freitas JK, Okur M, Khadka P, Kulikowicz T, Holland PP, Tian J, Croteau DL, Davis AJ et al (2017) Cell cycle-dependent phosphorylation regulates RECQL4 pathway choice and ubiquitination in DNA double-strand break repair. Nat Commun 8: 2039

Lu L, Harutyunyan K, Jin W, Wu J, Yang T, Chen Y, Joeng KS, Bae Y, Tao J, Dawson BC et al (2015) RECQL4 Regulates p53 Function In Vivo During Skeletogenesis. J Bone Miner Res 30: 1077–1089

Mahrour N, Redwine WB, Florens L, Swanson SK, Martin-Brown S, Bradford WD, Staehling-Hampton K, Washburn MP, Conaway RC, Conaway JW (2008) Characterization of Cullin-box sequences that direct recruitment of Cul2-Rbx1 and Cul5-Rbx2 modules to Elongin BC-based ubiquitin ligases. J Biol Chem 283: 8005–8013

Marino F, Mojumdar A, Zucchelli C, Bhardwaj A, Buratti E, Vindigni A, Musco G, Onesti S (2016) Structural and biochemical characterization of an RNA/DNA binding motif in the N-terminal domain of RecQ4 helicases. Sci Rep 6: 21501

Matson JP, Cook J, 2020. MCM Chromatin flow cytometry for cell cycle V.1, protocolsio. Springer Nature.

Matson JP, Dumitru R, Coryell P, Baxley RM, Chen W, Twaroski K, Webber BR, Tolar J, Bielinsky AK, Purvis JE et al (2017) Rapid DNA replication origin licensing protects stem cell pluripotency. Elife 6

Minocherhomji S, Ying S, Bjerregaard VA, Bursomanno S, Aleliunaite A, Wu W, Mankouri HW, Shen H, Liu Y, Hickson ID (2015) Replication stress activates DNA repair synthesis in mitosis. Nature 528: 286–290

Ng AJ, Walia MK, Smeets MF, Mutsaers AJ, Sims NA, Purton LE, Walsh NC, Martin TJ, Walkley CR (2015) The DNA helicase Recql4 is required for normal osteoblast expansion and osteosarcoma formation. PLoS Genet 11: e1005160

Padayachy L, Ntallis SG, Halazonetis TD (2024) RECQL4 is not critical for firing of human DNA replication origins. Sci Rep 14: 7708

Petkovic M, Dietschy T, Freire R, Jiao R, Stagljar I (2005) The human Rothmund-Thomson syndrome gene product, RECQL4, localizes to distinct nuclear foci that coincide with proteins involved in the maintenance of genome stability. J Cell Sci 118: 4261–4269

Pilcher C, Buco PAV, Truong JQ, Ramsland PA, Smeets MF, Walkley CR, Holien JK (2025) Characteristics of the Kelch domain containing (KLHDC) subfamily and relationships with diseases. FEBS Lett

Rusnac DV, Lin HC, Canzani D, Tien KX, Hinds TR, Tsue AF, Bush MF, Yen HS, Zheng N (2018) Recognition of the Diglycine C-End Degron by CRL2(KLHDC2) Ubiquitin Ligase. Mol Cell 72: 813–822 e814

Sangrithi MN, Bernal JA, Madine M, Philpott A, Lee J, Dunphy WG, Venkitaraman AR (2005) Initiation of DNA replication requires the RECQL4 protein mutated in Rothmund-Thomson syndrome. Cell 121: 887–898

Sanjana NE, Shalem O, Zhang F (2014) Improved vectors and genome-wide libraries for CRISPR screening. Nat Methods 11: 783–784

Scott DC, Chittori S, Purser N, King MT, Maiwald SA, Churion K, Nourse A, Lee C, Paulo JA, Miller DJ et al (2024a) Structural basis for C-degron selectivity across KLHDCX family E3 ubiquitin ligases. Nat Commun 15: 9899

Scott DC, Dharuman S, Griffith E, Chai SC, Ronnebaum J, King MT, Tangallapally R, Lee C, Gee CT, Yang L et al (2024b) Principles of paralog-specific targeted protein degradation engaging the C-degron E3 KLHDC2. Nat Commun 15: 8829

Singbrant S, Russell MR, Jovic T, Liddicoat B, Izon DJ, Purton LE, Sims NA, Martin TJ, Sankaran VG, Walkley CR (2011) Erythropoietin couples erythropoiesis, B-lymphopoiesis, and bone homeostasis within the bone marrow microenvironment. Blood 117: 5631–5642

Singh DK, Karmakar P, Aamann M, Schurman SH, May A, Croteau DL, Burks L, Plon SE, Bohr VA (2010) The involvement of human RECQL4 in DNA double-strand break repair. Aging Cell 9: 358–371

Smeets MF, DeLuca E, Wall M, Quach JM, Chalk AM, Deans AJ, Heierhorst J, Purton LE, Izon DJ, Walkley CR (2014) The Rothmund-Thomson syndrome helicase RECQL4 is essential for hematopoiesis. J Clin Invest 124: 3551–3565

Terui R, Berger SE, Sambel LA, Song D, Chistol G (2024) Single-molecule imaging reveals the mechanism of bidirectional replication initiation in metazoa. Cell 187: 3992–4009 e3925

Thakur BL, Redon CE, Fu H, Sebastian R, Kusi NA, Zhuang SZ, Pongor LS, Bohr VA, Aladjem MI (2025) Selective interactions at pre-replication complexes categorize baseline and dormant origins. Nat Commun 16: 4140

Timms RT, Koren I (2020) Tying up loose ends: the N-degron and C-degron pathways of protein degradation. Biochem Soc Trans 48: 1557–1567

Timms RT, Mena EL, Leng Y, Li MZ, Tchasovnikarova IA, Koren I, Elledge SJ (2023) Defining E3 ligase-substrate relationships through multiplex CRISPR screening. Nat Cell Biol 25: 1535–1545

Vignon C, Debeissat C, Georget MT, Bouscary D, Gyan E, Rosset P, Herault O (2013) Flow cytometric quantification of all phases of the cell cycle and apoptosis in a two-color fluorescence plot. PLoS One 8: e68425

Wang GG, Calvo KR, Pasillas MP, Sykes DB, Hacker H, Kamps MP (2006) Quantitative production of macrophages or neutrophils ex vivo using conditional Hoxb8. Nat Methods 3: 287–293

Wang LL, Gannavarapu A, Kozinetz CA, Levy ML, Lewis RA, Chintagumpala MM, Ruiz-Maldanado R, Contreras-Ruiz J, Cunniff C, Erickson RP et al (2003) Association between osteosarcoma and deleterious mutations in the RECQL4 gene in Rothmund-Thomson syndrome. J Natl Cancer Inst 95: 669–674

Wang LL, Levy ML, Lewis RA, Chintagumpala MM, Lev D, Rogers M, Plon SE (2001) Clinical manifestations in a cohort of 41 Rothmund-Thomson syndrome patients. Am J Med Genet 102: 11–17

Wang LL, Worley K, Gannavarapu A, Chintagumpala MM, Levy ML, Plon SE (2002) Intron-size constraint as a mutational mechanism in Rothmund-Thomson syndrome. Am J Hum Genet 71: 165–167

Wickham H (2016) ggplot2: Elegant Graphics for Data Analysis. Springer-Verlag New York

Wu J, Capp C, Feng L, Hsieh TS (2008) Drosophila homologue of the Rothmund-Thomson syndrome gene: essential function in DNA replication during development. Dev Biol 323: 130–142

Xu JJ, Chalk AM, Nikolic I, Simpson KJ, Smeets MF, Walkley CR (2022) Genome-wide screening identifies cell-cycle control as a synthetic lethal pathway with SRSF2P95H mutation. Blood Adv 6: 2092–2106

Yeh CW, Huang WC, Hsu PH, Yeh KH, Wang LC, Hsu PW, Lin HC, Chen YN, Chen SC, Yeang CH et al (2021) The C-degron pathway eliminates mislocalized proteins and products of deubiquitinating enzymes. EMBO J 40: e105846

Yen HC, Elledge SJ (2008) Identification of SCF ubiquitin ligase substrates by global protein stability profiling. Science 322: 923–929

Yen HC, Xu Q, Chou DM, Zhao Z, Elledge SJ (2008) Global protein stability profiling in mammalian cells. Science 322: 918–923

Zhang Z, Sie B, Chang A, Leng Y, Nardone C, Timms RT, Elledge SJ (2023) Elucidation of E3 ubiquitin ligase specificity through proteome-wide internal degron mapping. Mol Cell 83: 4191–4192

